# Glioblastoma cell fate is differentially regulated by the microenvironments of the tumour bulk and infiltrative margin

**DOI:** 10.1101/2021.06.11.447915

**Authors:** Claudia Garcia-Diaz, Elisabetta Mereu, Melanie P. Clements, Anni Pöysti, Felipe Galvez-Cancino, Simon P. Castillo, Lilas Courtot, Sara Ruiz, Federico Roncaroli, Yinyin Yuan, Sergio A. Quezada, Holger Heyn, Simona Parrinello

**Author notes:** these two authors contributed equally to the work (Claudia Garcia-Diaz, Elisabetta Mereu).

## Abstract

Glioblastoma recurrence originates from invasive cells at the tumour margin that escape surgical debulking, but their biology remains poorly understood. Here we generated three somatic mouse models recapitulating the main glioblastoma driver mutations to characterise margin cells. We find that, regardless of genetics, tumours converge on a common set of neural-like cellular states. However, bulk and margin display distinct neurogenic patterns and immune microenvironments. The margin is immune-cold and preferentially follows developmental-like trajectories to produce astrocyte-like cells. In contrast, injury-like programmes dominate in the bulk, are associated with immune infiltration and generate lowly-proliferative injured neural progenitor-like (iNPCs) cells. *In vivo* label-retention approaches further demonstrate that iNPCs account for a significant proportion of dormant glioblastoma cells and are induced by interferon signalling within T-cell niches. These findings indicate that tumour region is a major determinant of glioblastoma cell fate and therapeutic vulnerabilities identified in bulk may not extend to the margin residuum.

## Introduction

Glioblastoma (GBM) is the most common and aggressive primary brain tumour (Weathers and Gilbert, 2014). Current standard of care, consisting of maximally safe surgical resection followed by chemo- and radiotherapy remains ineffective, leading to invariable recurrence and a median survival of less than 18 months (Stupp et al., 2005).

A main cause of therapy-resistance is the ability of GBM cells to diffusely infiltrate into the normal brain (Cuddapah et al., 2014; Vehlow and Cordes, 2013). Infiltration precludes curative surgery, leading to tumour regrowth from cells that have invaded past the resection margin (Cuddapah et al., 2014; Vehlow and Cordes, 2013). Despite its crucial role in recurrence however, the invasive GBM margin remains poorly characterised (Vehlow and Cordes, 2013). This gap in our knowledge is largely due to the paucity of available patient material from the tumour margin, particularly from distal regions. Indeed, current knowledge of GBM biology originates almost exclusively from analysis of the tumour bulk collected during biopsy or surgical de-bulking. Nonetheless, as invasive cells, rather than bulk cells, give rise to recurrence, potential differences between bulk and margin tumour cells would have profound therapeutic implications.

The pervasive molecular and cellular heterogeneity of GBM further underlies recurrence by limiting efficacy of both standard and targeted therapies (Qazi et al., 2017). Large scale research efforts have carried out detailed molecular characterisation of human GBM, revealing marked genetic, epigenetic and transcriptional inter-tumoural heterogeneity (Brennan et al., 2013; Network, 2008; Sturm et al., 2012; Verhaak et al., 2010). Based on these analyses, GBMs have been classified into three main molecular subtypes, termed proneural, classical and mesenchymal, defined by distinct transcriptional signatures and associated driver mutations in PDGFRA, EGFR and NF1, respectively (Verhaak et al., 2010; Wang et al., 2017). In addition, GBMs display remarkable intra-tumour heterogeneity, with individual tumours containing co- existing cell populations of different genetics and subtypes (Couturier et al., 2020; Neftel et al., 2019; Patel et al., 2014).

At the cellular level, GBMs recapitulate developmental-like lineage hierarchies (Couturier et al., 2020; Lan et al., 2017; Neftel et al., 2019). The apex of this hierarchy is occupied by glioma stem-like cells (GSCs), defined by their ability to self-renew and differentiate into non-stem tumour cells (Galli et al., 2004; Lan et al., 2017; Lathia et al., 2015; Singh et al., 2004). GSCs are thought to play a key role in recurrence due to their tumour-initiation potential and intrinsic resistance to chemotherapy and radiation (Bao et al., 2006; Chen et al., 2012). Interestingly, GSCs were also shown to be more invasive than non-stem tumour cells (Cheng et al., 2011). This has led to the speculation that GSCs may drive infiltration at the invasive niche, but as other studies suggest a possible loss of stemness at the margin, their exact role in invasion remains an open question (Hoelzinger et al., 2005; Molina et al., 2010; Piccirillo et al., 2009). In analogy to neural development and adult neurogenesis, GSCs are also thought to be slow-cycling and give rise to actively dividing progenitor-like cells that in turn generate partially differentiated progeny (Obernier and Alvarez-Buylla, 2019). Within the tumour bulk, lineage progression occurs towards glia-like fate, including OPC-, astrocyte- and neural progenitor-like states, or to a mesenchymal-like phenotype (Couturier et al., 2020; Neftel et al., 2019). These state transitions are modulated by driver mutations and by the immune microenvironment, with common GBM mutations biasing towards either OPC (PDGFR) or astrocyte-like fate (EGFR), and NF1-dependent high microglia/macrophage infiltration promoting a mesenchymal-like fate (Hara et al., 2021; Neftel et al., 2019; Wang et al., 2017). In contrast, little is known about the lineage progression of invasive cells. Yet, the microenvironments of the bulk and margin are dramatically different, with the bulk comprising hypoxic, necrotic and angiogenic regions and the margin containing largely normal brain tissue (Brooks and Parrinello, 2017). This suggests that distinct pressures on tumour cell fate choice might exist between the two regions and that knowledge of bulk heterogeneity may not directly inform margin phenotypes.

Here, we investigated the biology of invasive GBM cells and how they are affected by genetic heterogeneity. We developed three somatic mouse models of GBM that carry the main subtype-associated patient mutations and share remarkable similarities with the human disease. By labelling the tumour cells with fluorescent reporters and exploiting the full accessibility of the murine invasive front, we used these models to compare bulk and margin tumour cells by single cell RNA-sequencing (scRNA-seq) and in functional studies. We found that regardless of underlying mutations, all three models converge on a finite set of cellular states that resemble normal neural cell types. However, the cancer hierarchy is distinctly modulated by tumour region. In the bulk, injury-like neurogenic programmes are dominant. This correlates with selective immune infiltration of the bulk and results in the generation of slow-cycling cells with properties of injured neural progenitor cells (iNPCs). In contrast, neurodevelopmental-like hierarchies biased towards astrocyte-like differentiation are prevalent at the margin, where the immune microenvironment resembles that of normal brain. We also show that the injured NPC state represents a large proportion of the GBM dormant population and is induced by high interferon signalling from T-cells that form bulk-specific niches. Our work reveals striking differences between bulk and margin biology and suggests that tumour region is a major determinant of GBM fate.

## Results

### Development of somatic GBM mouse models

To characterise the biology of invasive tumour cells and how it is affected by genetic alterations, we developed three somatic mouse models of GBM carrying combinations of mutations commonly associated with the main human subtypes (Network, 2008; Verhaak et al., 2010; Wang et al., 2017). This enabled us to directly link tumour phenotypes to disease-relevant driver mutations and model the heterogeneity of human GBM through combined analysis of the three mouse models. Furthermore, the introduction of a tdTomato reporter in all tumour cells allowed us to comprehensively sample the tumour margin and discriminate tumour cells from normal brain cells based on tdTomato fluorescence.

Subtype-relevant mutations were introduced into endogenous neural stem cells (NSCs) of the subventricular zone neurogenic niche (SVZ), a frequent cell of origin in GBM patients (Alcantara Llaguno et al., 2009; Lee et al., 2018). Specifically, the following mutations were used: EGFRvIII overexpression and *Cdkn2a* knock-out (hereon EGFR model); *Pdgfra* overexpression and *Trp53* knock-out (hereon Pdfgra model); *Nf1, Pten* and *Trp53* knockout (hereon Nf1 model). To this end, a non-integrating plasmid encoding for the PiggyBase transposase alone (Pdgfra and EGFR models) or with Cas9 (Nf1 model) together with an integrating piggyBac vector carrying the oncogenes, CRISPR guides to tumour suppressors, Cre recombinase and tdTomato were co-electroporated into the lateral ventricles of *Trp5^fl/fl^* (Pdgfra and Nf1 models) or *Cdkn2a^fl/fl^* (EGFR model) of P2 pups (Figure 1A) (Chen and LoTurco, 2012). Upon electroporation, transient Cas9/gRNA expression results in inactivation of the tumour suppressor genes, whereas PiggyBase-mediated integration of the piggyBac vector ensures stable expression of the oncogenes and the td-Tomato reporter in the targeted NSCs and their progeny. To ensure selective targeting of neural stem cells (NSC), Cas9 and Cre expression were driven by a truncated version of the human GFAP promoter (herein hGFAPMIN) previously reported to maintain the specificity of the full GFAP promoter while increasing its activity (Lee et al., 2008). Promoter specificity was confirmed by electroporation of a hGFAPMIN-tdTomato reporter construct, which revealed selective tomato expression in NSCs with radial glia morphology that were largely Ki67^-^/GFAP^+^ (Supplemental Figure 1A-C). All genotypes generated tdTomato^+^ tumours with histological and molecular features of GBM, including vascular proliferation and necrosis, as well as expression of the GBM markers Sox2, Olig2 and GFAP, within 8-15 weeks and with high penetrance (Figure 1B-E). Western analysis of primary cells acutely isolated from the three tumour types confirmed that the mutations were correctly introduced in each model (Supplemental Figure 1D). Together, these results suggest that the models closely recapitulate the human disease and can inform on GBM biology.

**Figure 1.**
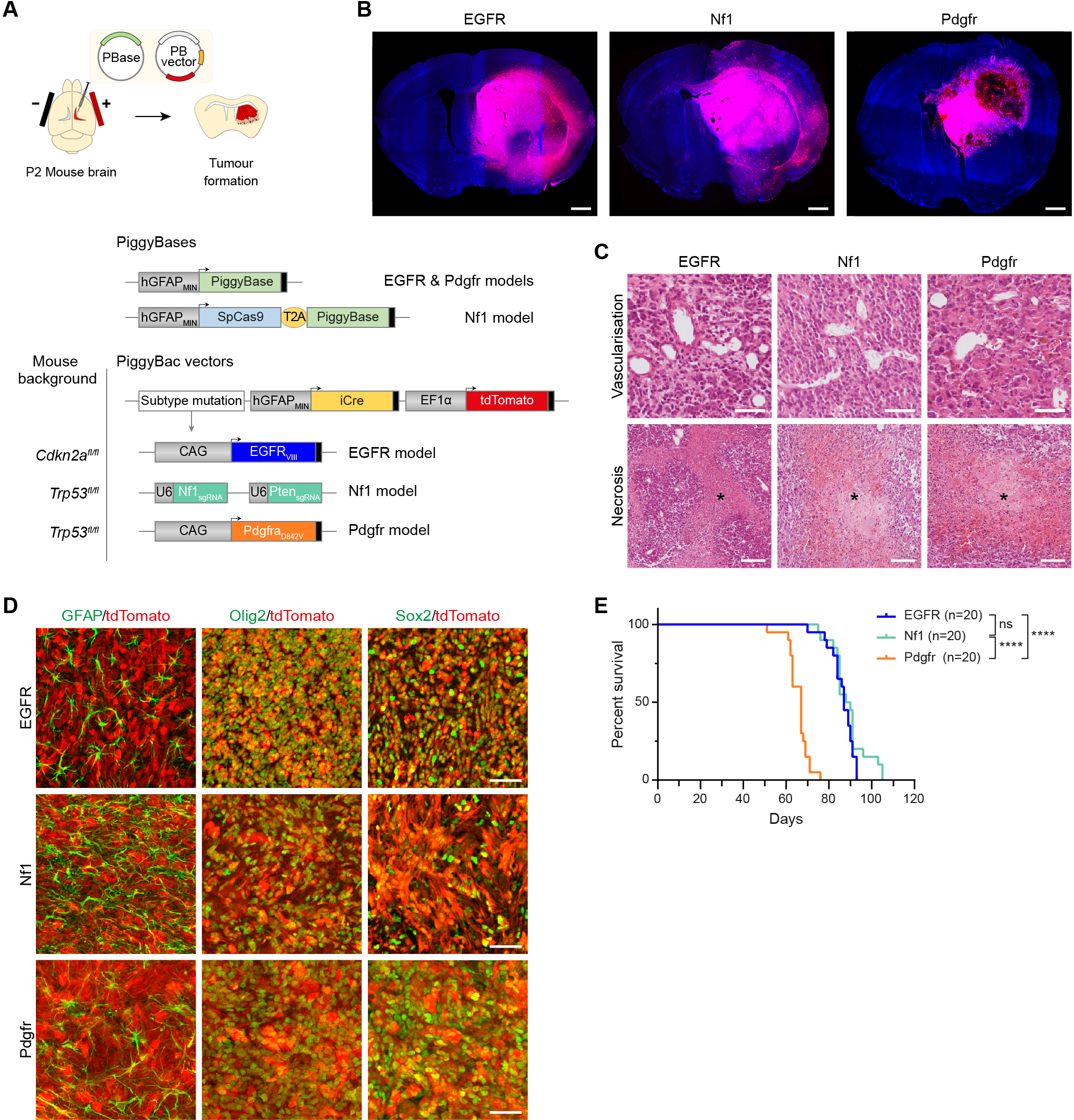
Development of somatic mouse models of GBM. **A**, Schematic of method for tumour generation and piggyBac constructs. **B**, Representative tile scan fluorescence images of tumours of each genotype. Tumour cells are labelled by endogenous tdTomato expression (red). Sections were counterstained with DAPI (blue). Scale Bar=1mm. **C**, Representative haematoxylin-eosin (H&E) stainings of tumour models showing examples of microvascular proliferation (top) and necrosis (asterisks, bottom). Scale Bars=50µm and 100µm, respectively. **D**, Immunofluorescence staining for the GBM markers GFAP, Olig2 and Sox2 (green) of tdTomato^+^ (red) EGFR, Nf1 and Pdgfr tumours, as indicated. Scale Bar=50µm. **E**, Kaplan- Meier survival plots of the three GBM models. n=20. Log Rank Mantel Cox test. (ns: p=0.1583; ****: p<0.0001). Median survival of 87, 89 and 67 days for EGFR, Nf1 and Pdgfr models, respectively. See also Supplemental Figure 1.

### Tumour cell states differ between bulk and margin

We next used scRNA-seq to profile invasive tumour cells and their bulk counterparts in each model. The bulk and striatal margin regions of three tumours of each genotype were microdissected under fluorescence guidance, enzymatically dissociated to single cells and FACS-sorted based on tdTomato fluorescence (Figure 2A and Supplemental Figure 2A) (Brooks et al., 2021). As expected, the proportion of tdTomato^+^ tumour cells was lower at the margin relative to the bulk, confirming accuracy of microdissection (Supplemental Figure 2B). Transcriptomes of an average of 470 cells (ranging from 410 to 531) per region were analysed using SMART-seq2 protocols (Figure 2A) (Picelli et al., 2014).

**Figure 2.**
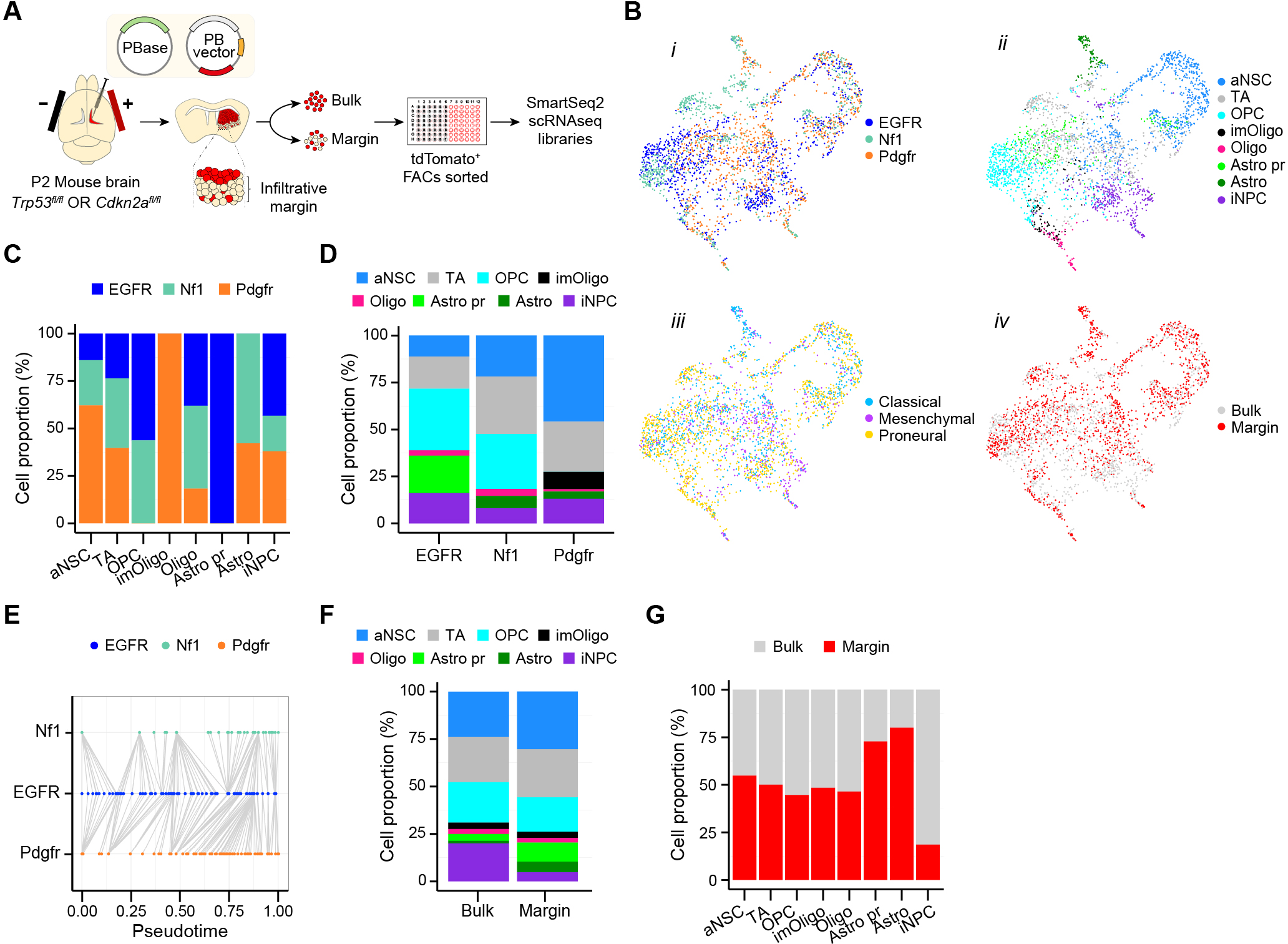
Tumour cell states differ between bulk and margin. **A**, Schematic of experimental set up. **B**, UMAP visualisation of 2824 cells from the combined GBM tumour models. Cells are coloured by *i*) genotype, *ii*) cell type, *iii*) Verhaak molecular subtyping and *iv*) tumor region (bulk, margin). **C**, Relative frequency of cell types across tumour genotypes. **D**, Cell type composition of the three models. **E**, Schematic representation of pseudotime alignment of differentiation trajectories in EGFR, Pdgfr and Nf1 tumours. The EGFR model was used as a reference for comparison with the other two models. **F**, Cell type composition of bulk and margin regions in the combined tumour models. **G**, Relative proportions of cell types in the bulk and margin regions of the combined tumour dataset. See also Supplemental Figure 2.

We first assessed the cellular composition of the tumours irrespective of tumour region. Each tumour model was first analysed independently and then all datasets were combined to identify common transcriptional patterns across the three genotypes. Data integration, based on canonical correlation analysis, revealed that cells did not segregate by genotype, but rather intermixed, converging onto 8 main subpopulations or states across regions (Figure 2B*i,* 2B*ii* and 2B*iv*). This is consistent with previous findings in human GBM (Couturier et al., 2020; Neftel et al., 2019) and is indicative of common and mutation-independent biological processes. Furthermore, all tumours contained mixtures of cells of all three transcriptional subtypes and four cellular states identified in patients, further confirming the validity of our models (Figure 2B*iii* and Supplemental Figure 2C) (Neftel et al., 2019; Wang et al., 2017).

As our models derive from the transformation of normal postnatal NSCs, we reasoned that the 8 identified clusters might correspond to states that mirror SVZ neurogenesis. We therefore compared expression signatures in each cluster with published scRNA-seq analyses of NSCs and their progeny in the normal and ischemic SVZ, which we hypothesized may be a state relevant to tumourigenesis due to the known links between injury and cancer (Supplemental Figure 2D-G and Supplemental Table 1) (Dvorak, 1986; Kalamakis et al., 2019; Llorens-Bobadilla et al., 2015; Mizrak et al., 2019). All tumours contained cells with signatures of normal or injured neural progenitors (Figure 2B*ii* and Supplemental Figure 2D-H), 4 of which were shared among all genotypes. Specifically, all tumours contained cells similar to active NSCs (aNSC), transit amplifying progenitors/early neuroblasts (TA), oligodendrocytes (oligo) and injured NPCs that result from ischemic brain injury (iNPC). The iNPC state included, but was not restricted to mesenchymal-like cells described by Neftel et al. In addition, EGFR and Nf1 tumours contained cells with signatures of oligodendrocyte precursor cells (OPCs) and Pdgfr and Nf1 tumours cells with astrocyte-like subpopulations (Figure 2C, D). Interestingly, although Pdgfr tumours lacked OPCs, they uniquely contained a subpopulation of cells with signatures of immature oligodendrocytes (imOligo), indicating that *Pdgfra* overexpression promotes maturation down the oligodendrocyte lineage, while concomitantly preventing further differentiation to more mature oligodendrocytes, in line with its developmental roles (Figure 2C, D) (Brooks et al., 2021; Zhu et al., 2014). Similarly, although EGFR tumours lacked more mature astrocyte-like cells, they contained a subpopulation with signatures of astrocyte progenitor-like cells (Astro pr), which was absent in the other two genotypes (Figure 2C, D). This suggests that EGFRvIII overexpression biases tumour cells towards astrogliogenesis, as previously reported for wildtype EGFR, while again preventing full differentiation (Neftel et al., 2019). Consistent with a differentiation block caused by mutant RTK signalling, pseudotemporal alignment of the differentiation trajectories of the three models revealed that both EGFR and Pdgfr tumours appeared more immature than Nf1 tumours lacking constitutive RTK activity, with EGFR tumours being the most immature (Figure 2E). Thus, our models indicate that regardless of genetics, tumour fates converge on a finite set of phenotypes that mimic neurogenesis, with driver mutations biasing towards specific cell fates, as observed in human GBM (Neftel et al., 2019). They also reveal that mutations control the extent by which tumour cells differentiate, with sustained developmental RTK signalling blocking lineage progression at immature progenitor-like states.

Next, we examined the impact of tumour region on cellular states, by comparing the frequency of the identified clusters in the bulk and margin of the tumours (Figure 2B*iv*, F, G). We found that while all cell fates were detected in both regions, location influenced the frequency of specific cell states, with the bulk being enriched for iNPCs and the margin for astrocyte-like fate. Interestingly, these biases were largely independent of genetics, as they were observed in all models, regardless of basal mutation-dependent lineage bias (Supplemental Figure 2I). These findings suggest that tumour region is dominant over driver mutations in modulating cell state and that margin and bulk biology differ.

### aNSC-like cells sit at the top of the tumour hierarchy and are not enriched at the margin

To better understand the tumour hierarchy in our models, we performed pseudotime analysis of the integrated datasets using normal SVZ neurogenesis trajectories to infer directionality (Figure 3A and Supplemental Figure 3A) (Mizrak et al., 2019). We found that the aNSC compartment, which was the most highly proliferative tumour subpopulation, was at the apex of the tumour hierarchy (Figure 3B and Supplemental Figure 3B). In analogy to findings in human GBM, this suggests that the aNSC subpopulation corresponds to GSCs within our models (Couturier et al., 2020). GSCs have been hypothesized to drive invasion based on the observation that GSCs-derived tumours are more invasive than tumours derived from non-stem tumour cells (Cheng et al., 2011). However, analyses of the stemness potential of invasive cells *in vivo* remain inconclusive, largely due to the challenges associated with profiling the human GBM margin (Hoelzinger et al., 2005; Molina et al., 2010; Piccirillo et al., 2009). We therefore took advantage of the accessibility of the margin in our models to explore the role of tumour stem-like cells in invasion and in the context of driver mutations.

**Figure 3.**
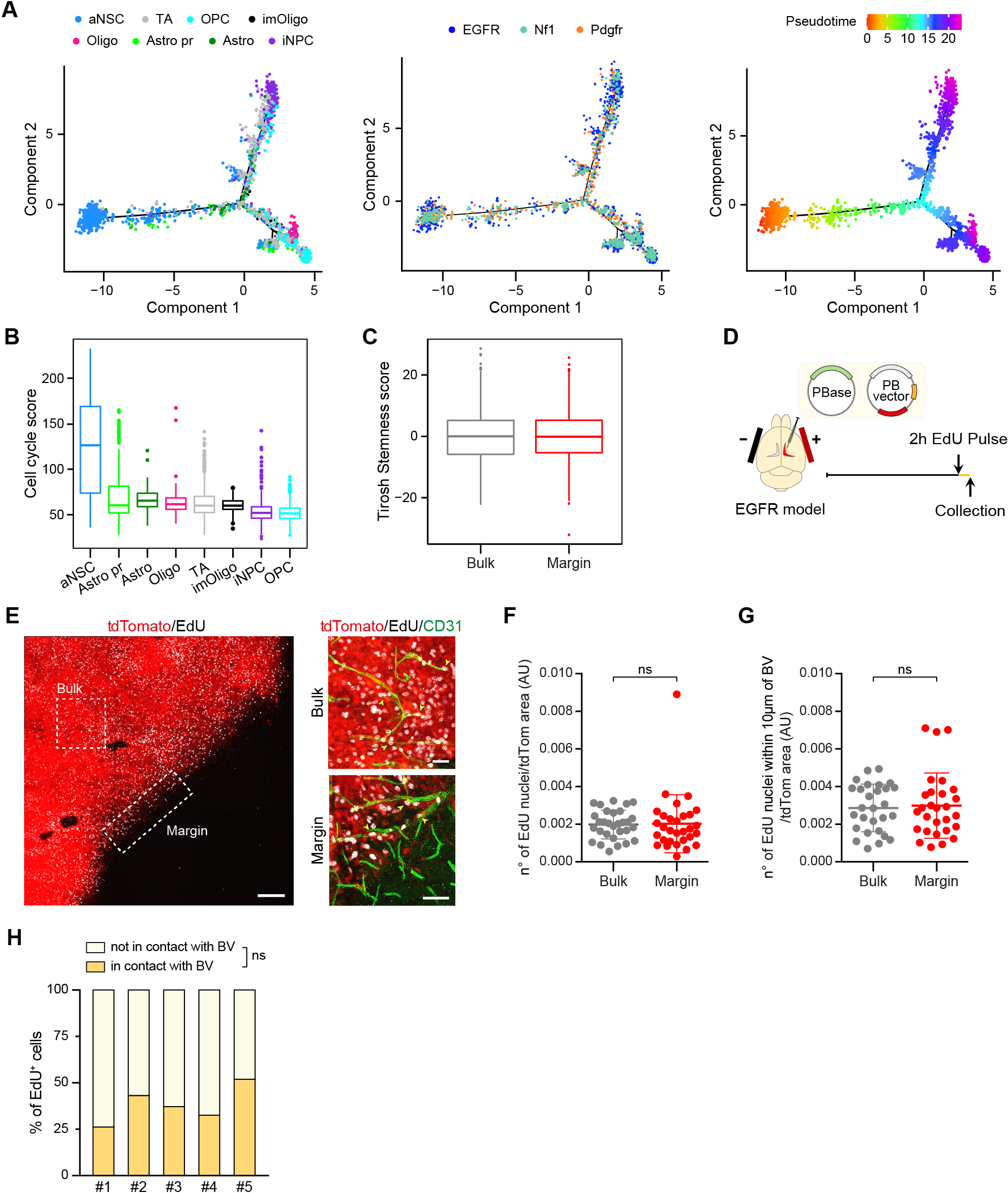
aNSC-like cells sit at the top of the tumour hierarchy and are not enriched at the margin. **A**, Differentiation trajectory analysis of 2824 GBM cells coloured by (from left to right): cell type, genotype and pseudotime by which cells are ordered along the differentiation path. **B**, Boxplots comparing the cell-cycle signature score in each tumour cell population within the combined tumour dataset. Boxplots are ordered from higher to lower cell cycle score and display the minimum, 1st, 2nd, 3rd quartile and maximum of scores for each cell type. **C**, Boxplots comparing the Tirosh stemness signature score in bulk and margin tumor regions. Boxplots display the minimum, 1st, 2nd, 3rd quartile and maximum of scores for each regions. **D**, Schematic of experimental set up. **E**, Representative immunofluorescence staining for EdU (grey) and the vascular marker Cd31 (green) of tdTomato^+^ EGFR tumours. Scale bars=200 µm and 50µm for close-up regions. **F**, Quantifications of the number of EdU^+^ GSCs in the bulk (grey) and margin (red) of EGFR tumours shown in E. n=5 tumours. Six regions of interest (ROIs) per tumour were counted. Two-tailed Mann-Whitney test (ns: p=0.4581). **G**, Quantifications of the number of EdU^+^ aNSC-like cells in the perivascular region within the bulk (grey) and margin (red) of EGFR tumours shown in E. n=6 tumours. Three ROIs per tumour were counted. Two-tailed Mann-Whitney test (ns: p=0.7389). **H**, Quantification of proportion of margin EdU^+^ aNSC-like cells with or without association with the invasive vasculature. n=5 tumours. Each independent repeat is plotted. Six ROIs per tumour were counted. Two-tailed paired Student’s t-test (ns: p=0.0567). See also Supplemental Figure 3.

Consistent with an equal distribution of aNSCs in bulk and margin (Figure 2G), we found no changes in stemness signatures between the two regions, suggesting that stem cell potential may not be a prerequisite for invasion (Figure 3C) (Tirosh et al., 2016). To test this more directly, we experimentally validated the distribution of aNSCs within tumours by immunofluorescence analysis using the EGFR model as a paradigm. We chose EGFR tumours for this and all later validation experiments because they reflected all phenotypes shared across the models, while displaying the simplest hierarchical organisation (Supplemental Figure 3A). As aNSC-like tumour cells were the most proliferative cells amongst all tumour cell types, we used their cell cycle characteristics to label them selectively within tumour sections, as previously reported for SVZ neurogenesis (Supplemental Figure 3C) (Codega et al., 2014; Ponti et al., 2013). Mice were given a 2h EdU pulse prior to sacrifice and terminal tumours were analysed for distribution of EdU^+^ aNSCs (Figure 3D). aNSC-like cells were evenly distributed across the entire tumour mass, as shown by quantification of EdU^+^ cells relative to tdTomato signal in both regions (Figure 3E, F). EdU^+^ cells were also not enriched within the perivascular space of the margin, one of the main invasive niches for GBM (Figure 3G, H). Indeed, a larger proportion of EdU^−^ tdTomato^+^ tumour cells than EdU^+^ cells invaded perivascularly (Supplemental Figure 3D). Thus, aNSC-like cells and more committed tumour progenitors appear to have comparable invasive potential, suggesting that invasion is driven by all tumour compartments.

### Differential evolution of cell states in bulk and margin

We next asked how the different cell states we identified in our models evolve from aNSCs and whether tumour hierarchies vary between bulk and margin. As shown in Figure 3A, pseudotime analysis was consistent with aNSCs undergoing lineage progression along two main routes, a developmental-like route that bifurcated to give rise to astrocyte-like or oligodendrocyte-like cells and an injury-like route that terminated with iNPC-like cells. Interestingly, when analysed in the context of tumour region, it became apparent that location impacted the tumour hierarchy in all models (Figure 4A and Supplemental Figure 4A). Progression to oligodendrocyte-like fate along the developmental route occurred in both bulk and margin, but astrocyte fate was favoured in the margin, consistent with the cell fate distributions measured in Figure 2G. In contrast, progression to the iNPC state along the injury-like route occurred almost exclusively in the bulk (Figure 4A and Supplemental Figure 4A). Consistent with a propensity for differentiation at the margin, invasive cells were overall more mature than bulk cells as judged by their global pseudotime alignment (Figure 4B).

**Figure 4.**
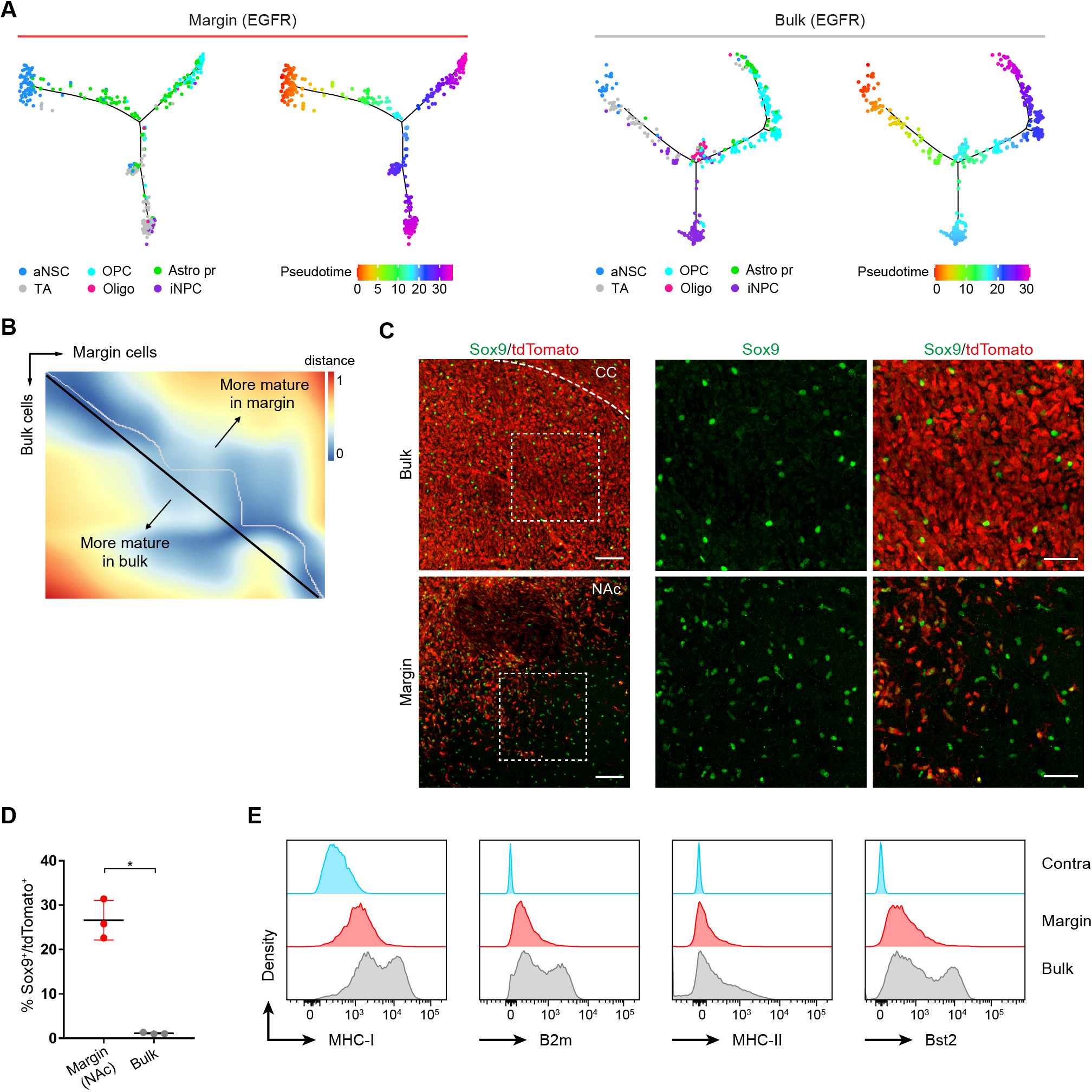
Differential evolution of cell states in bulk and margin. **A**, Differentiation trajectory analysis of 463 margin (left) and 494 bulk (right) EGFR cells coloured by cell type. **B**, Distance matrix representative of the global alignment of bulk and margin pseudotimes. Trajectory starts at the top left corner and ends at the bottom right corner of the plot. **C**, Representative immunofluorescence image of the bulk and margin regions of tdTomato^+^ EGFR tumours stained for Sox9 (green). Images on the right are magnifications of the boxed regions on left images. Scale bar=100µm and 50µm for close-up regions. CC: Corpus callosum, NAc: nucleus accumbens. **D**, Quantifications of the number of Sox9^+^ astrocyte-like cells in the tumours shown in D. n=3 EGFR tumours >150 cells per tumour were counted. Paired two-tailed Student’s t test. Mean±SD. **E**, FACS analysis of expression profiles for iNPC markers MHC-I, B2m, MHC-II and Bst2 in the bulk, margin and contralateral (Contra) regions of EGFR tumours. Representative histogram shown. n=3 tumours. See also Supplemental Figure 4.

To determine whether the cell state changes inferred from transcriptional signatures corresponded to phenotypic changes, we examined the distribution of astrocyte-like cells and iNPCs in EGFR tumours at the protein level and in their spatial context. We used Sox9, a master regulator of astrogliogenesis and one of the most differentially expressed genes in the astrocyte-like clusters (Supplemental Table 1), as a marker for Astro pr (Rowitch and Kriegstein, 2010). Immunofluorescence analysis of tumour sections confirmed that Sox9^+^ cells were rare within the bulk of the tumour and increased as cells invaded into the striatum (Figure 4C). However, Sox9 upregulation was heterogeneous within the margin, with a striking enrichment in the nucleus accumbens and a more modest increase across the dorsal striatum (Figure 4C). Quantification of the percentage of Sox9^+^/tdTomato^+^ cells within the nucleus accumbens revealed that as much as a quarter of all tumour cells acquired Astro pr fate in this region (Figure 4D). Thus, the margin microenvironment imposes distinct and regionally- determined selective pressures on tumour cells. To examine the distribution of iNPCs, we selected the MHC class I markers H-2K^b^ and β₂ microglobulin (B2m), Bst2 and MHC class II I-A/I-E as marker genes as they were all markedly increased in this cluster (Supplemental Figure 2H and Supplemental Table 1). The bulk and margin regions of 3 EGFR tumours were microdissected under fluorescence guidance, dissociated to single cells, immunolabelled and subjected to FACS analysis. In agreement with the bioinformatics data, we found a much greater proportion of MHC-I^high^, MHC class II^high^, B2M^high^ and Bst2^+^ tdTomato^+^ iNPCs in the bulk of the tumour relative to the margin, confirming that iNPC fate evolves selectively in the bulk (Figure 4E and Supplemental Figure 7). Thus, the tumour hierarchy is biased by location and subject to significant extrinsic control.

### iNPCs comprise a large proportion of dormant tumour cells

In the ischemic SVZ, injured neural progenitors include primed quiescent cells that are poised for activation (Llorens-Bobadilla et al., 2015). As this cellular compartment was also characterised by low proliferation in our models (Figure 3B), we hypothesised that iNPCs may represent dormant/quiescent tumour cells. To test this, we modified the EGFR piggyBac construct to incorporate a Tet-ON inducible H2B-GFP reporter of label retention (Foudi et al., 2009). This approach allows *in vivo* detection of slow-cycling tumour cells by pulse-chase experiments using doxycycline (Dox) administered in the drinking water. To simplify the piggyBac system and enable transformation of endogenous NSCs in any mouse genetic background, we also introduced gRNAs for *Cdkn2a* into the piggyBac backbone (Figure 5A), producing a fully integrated genetic tool for tumour initiation. Similar to the EGFR piggyBac system, the EGFR-H2B-GFP construct produced tumours with histological features of GBM and with high penetrance, but with shorter latency, likely due to more efficient integration of the modified vector (Figure 6F and Supplemental Figure 5A).

**Figure 5.**
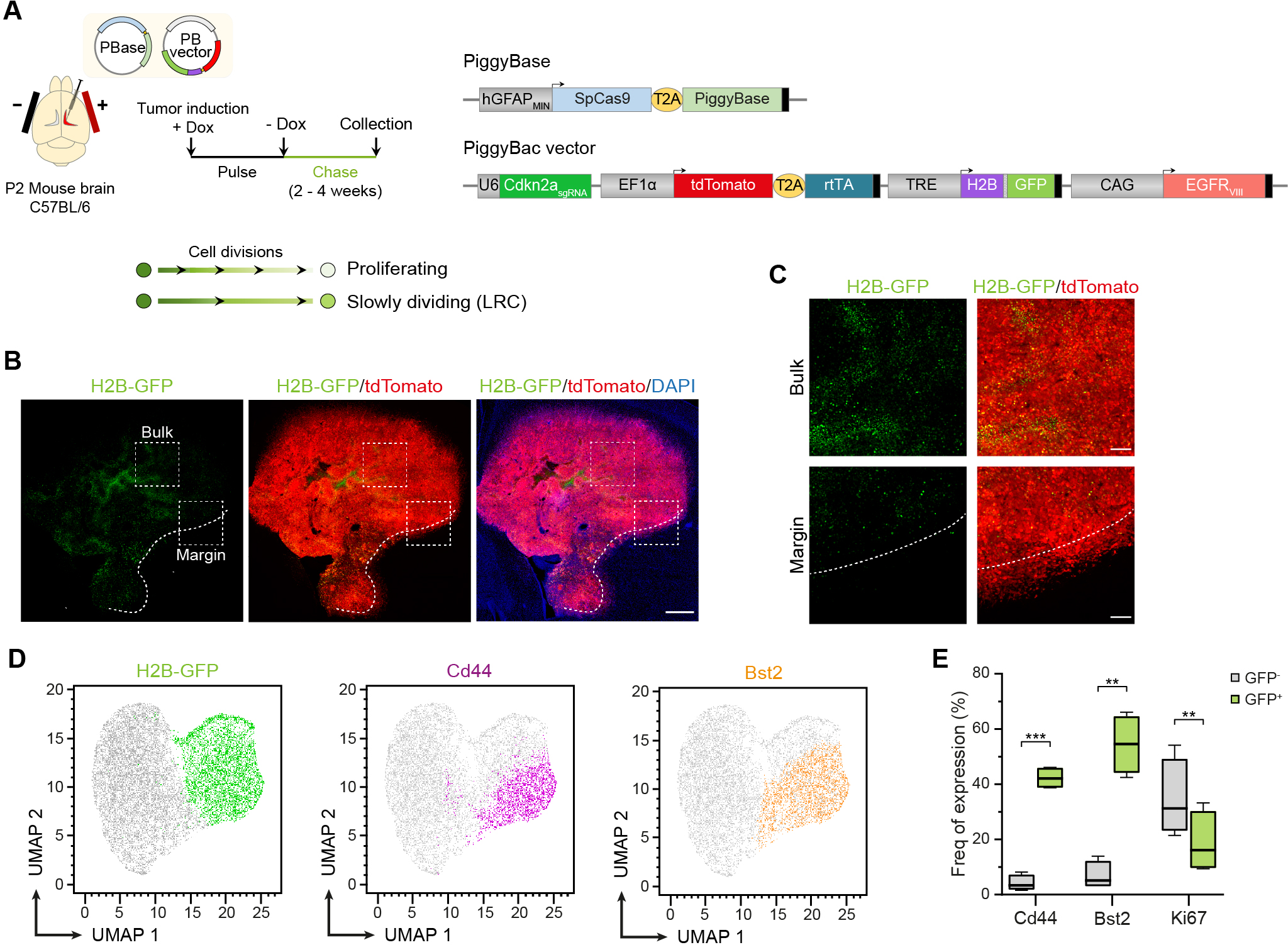
iNPCs comprise a large proportion of dormant tumour cells. **A**, Schematic representation of EGFR-H2B-GFP label-retention tumour model. Dox, doxycycline. LRC, label retaining cells. **B**, Immunofluorescence tile scan image of a representative EGFR-H2B- GFP tumour at 2 weeks of doxycycline chase. Endogenous tdTomato^+^ (red) and H2B-GFP^+^ (green) fluorescence is shown. Sections were counterstained with DAPI. Note that LRC are restricted to the tumour bulk. Scale bar=500µm. **C**, Higher magnification images of tumour boxed regions of the bulk and margin of the tumour shown in B. Scale bar=100µm. Dotted line demarcates margin. **D**, UMAP projections of expression of GFP and iNPC markers Cd44 and Bst2. Bulk tumour regions from four EGFR-H2B-GFP tumours at two weeks of doxycycline chase were analysed by FACs. 3,000 tumour cells (Cd45^-^ tdTomato^+^) were concatenated from each tumour and gated positive populations were projected onto UMAP. **E**, Quantification of the proportion of Bst2, Cd44 and Ki67-expressing cells H2B-GFP negative (GFP^-^) and GFP positive LRC tumour cells (GFP^+^) from experiment shown in D. Whisker plots show median and min-max value range. n=4. Two-tailed paired Student’s t-test (Cd44: p=0.0003, Bst2: p=0.0051 and Ki67: p=0.0047). See also Supplemental Figure 5.

**Figure 6.**
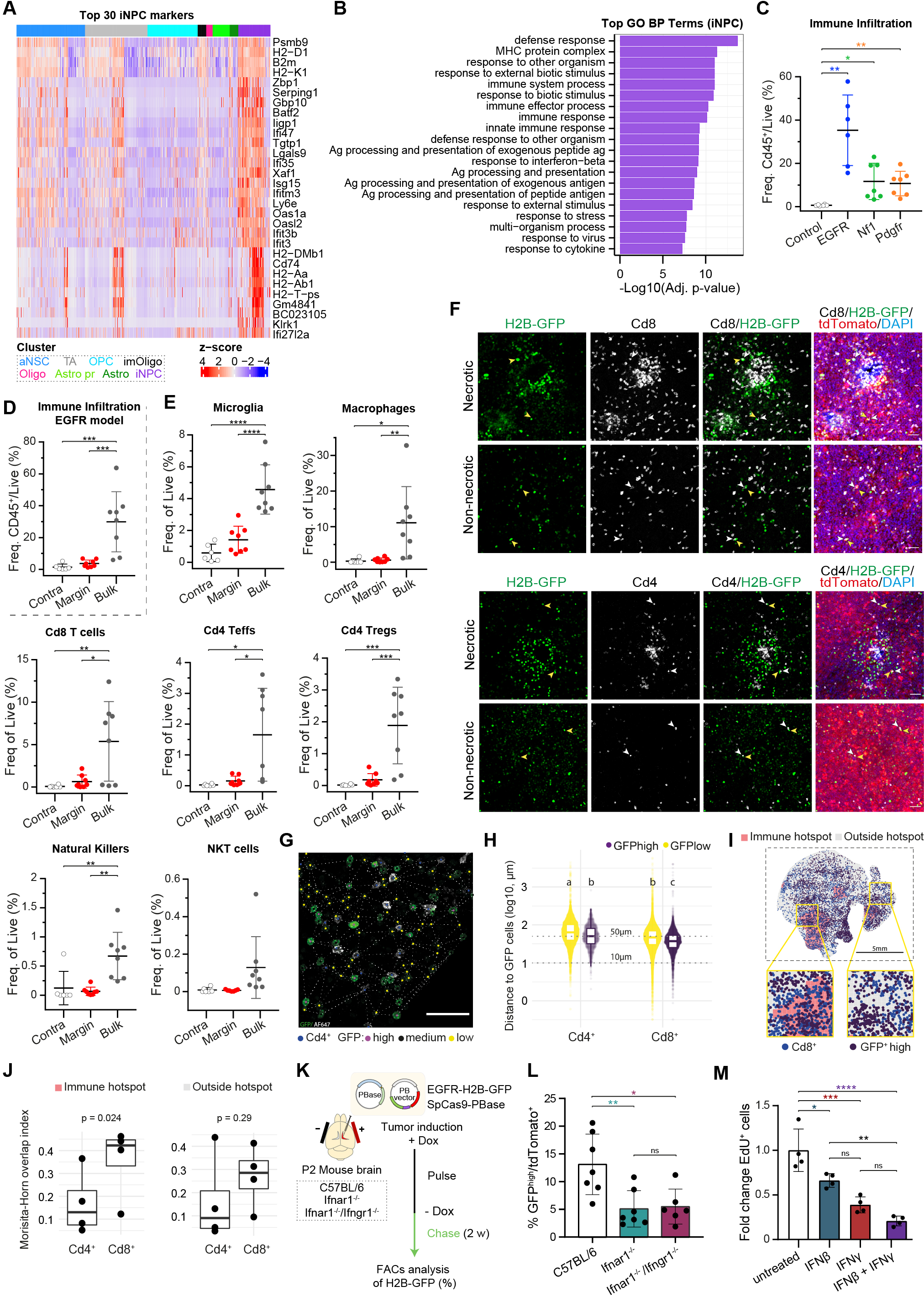
Dormant tumour cells are induced by interferon in T-cell niches. **A**, Heatmap of the top 30 markers of the iNPC population. Columns are grouped by cell type (top bar). Normalized gene expressed values (z-scores) are shown. **B**, Gene Ontology (GO) enrichment analysis for iNPCs from the integrated analysis of all genotypes. Shown are the top-20 GO terms ordered by log10 of their relative adjusted p-value. **C**, FACS analysis of the percentage of Cd45^+^ immune cells over total number of live cells in the three tumour models and normal control brains. n=6 for Control; n=6 for EGFR; n=7 for Nf1 and n=7 for Pdgfr. Welch and Brown-Forsythe one-way ANOVA with Dunnet’s T3 multiple comparison test. Control vs EGFR: p=0.0092; Control vs Nf1: p=0.0356; Control vs Pdgfr: p=0.0095. Mean±SD. **D**, FACS analysis of the percentage of Cd45^+^ immune cells over total number of live cells in the bulk and margin regions of EGFR tumours and contralateral non-infiltrated brain (Contra). n=6 for contralateral; n=8 for bulk and margin. One-way ANOVA with Tukey test. Mean±SD. **E**, FACS analysis of the percentage of indicated immune populations over total number of live cells in the bulk and margin regions of EGFR tumours and contralateral non-infiltrated brain (Contra). n=6 for contralateral; n=8 for bulk and margin. One-way ANOVA with Tukey test. Mean±SD. **F**, representative immunofluorescence staining for Cd8 (top) and Cd4 (bottom) T cells in the bulk of EGFR-H2B-GFP tumours. Shown are examples of T-cell rich necrotic regions surrounded by GFP positive LRCs (top) and direct cell-cell interactions between LRCs (yellow arrows) and T-cells (white arrows, bottom). Scale bars=100µm. **G**, Example of cell detection, classification, and spatial distance to neighbouring cells after Delaunay triangulation (segmented white lines). Scale bar=50µm **H**, Quantification of distances between H2B- GFPhigh (GFPhigh) or H2B-GFPlow (GFPlow) tumour cells and Cd4 or Cd8 T cells. Different letters above boxplots indicate statistical differences between groups quantified with a Tukey’s test after a significant generalised linear mixed model for hypothesis testing. **I**, Schematic representation of the distribution of Cd8 T cells and H2B-GFPhigh (GFPhigh) cells in an EGFR-H2B-GFP tumour. Regions of T cell hotspots, detected by Getis-Ord’s G* are shown in pink. **J**, Measurement of Morisita-Horn overlap indexes of indicated comparisons. An index higher than 0 indicates a non-random distribution of cells. p-value relates to comparison of H2B-GFPhigh (GFPhigh) cell co-localisation with Cd4 or Cd8 T cells. n=4 tumours, F test. **K**, Schematic representation of experimental set up. **L**, FACS quantification of percentage of H2B-GFP^high^ LRC in EGFR-H2B-GFP tumours generated in wildtype (WT), *Ifnar1^-/-^* or *Ifnar1^-/-^*;*Ifngr1^-/-^* animals. n=7 for WT, n=6 for *Ifnar1^-/-^*, n=7 for *Ifnar1^-/-^*;*Ifngr1^-/-^*. One-way ANOVA with Tukey’s multiple comparison test **: p= 0.0058; *: p=0.013; ns: p=0.9829. Mean±SD. **M**, FACS quantification of the percentage of EdU^+^ cells in cultured primary H2B-GFP^-^ tumour cells isolated from EGFR tumours and left untreated or treated with the indicated recombinant interferons for 48h. n=4 repeats. One-way ANOVA with Tukey’s multiple comparison test. Untreated vs IFN*β*: p=0.0194; untreated vs IFN*γ*: p=0.0002; untreated vs IFN*β*+IFNg: p<0.0001; IFN*β* vs IFN*γ*: p=0.0619; IFN*β* vs IFN*β*+IFN*γ*: p=0.0024; IFN*γ* vs IFN*β*+IFN*γ*: p=0.2872. Mean±SD. See also Supplemental Figure 6.

Dox was administered from electroporation until week 5 of tumour development followed by a 2- to 4-week chase period (Figure 5A). Immunofluorescence analysis before and after chase indicated that the H2B-GFP protein was efficiently incorporated in the chromatin of >90% of all tumour cells and effectively diluted over the chase period (Supplemental Figure 5B and C). Furthermore, by 2 weeks of chase, H2B-GFP^+^ cells were largely negative for the proliferation marker Ki67, which was instead restricted to H2B-GFP^-^ and a minority of H2B-GFP^low^ cells, confirming that the approach successfully identified dormant tumour cells (Supplemental Figure 5D). We next examined the distribution of label retaining H2B-GFP^+^ cells (LRC) within the tumour *in situ*. In agreement with the enrichment of iNPCs in the bulk identified by scRNA-seq (Figure 2G), immunofluorescence analysis revealed that the majority of LRC were found in the tumour bulk (Figure 5B, C). Interestingly, their distribution was not uniform, but rather restricted to specific bulk regions, with LRCs often found in clusters, suggestive of microenvironmental regulation (Figure 5B, C). In addition, FACS analysis indicated that H2B- GFP^+^ cells were selectively enriched for expression of the iNPC markers Cd44 and Bst2 relative to H2B-GFP^-^ tumour cells (Figure 5D, E and Supplemental Figure 7). We conclude that iNPCs are LR tumour cells, induced to enter a dormancy-like state within the bulk of the tumour.

### Dormant tumour cells are induced by interferon in T-cell niches

To understand the signals that induce dormancy in the tumour bulk, we examined in greater detail the gene expression profile of iNPCs. Analysis of top marker genes alongside gene ontology analysis showed an overrepresentation of immune genes and signatures, particularly those linked to interferon signalling, as reported in the ischemic SVZ (MHC class proteins, innate immune response, antigen processing, response to virus, response to interferon beta) (Figure 6A, B, Supplemental Figure 6A and Supplemental Table 2) (Llorens-Bobadilla et al., 2015). This suggested that the iNPC state may be induced by interactions with immune cells. To explore this idea, we examined the distribution of the main immune compartments in our tumour models. We found that all three models were infiltrated by immune cells as in the human disease (Pombo Antunes et al., 2020), with the EGFR model displaying a trend towards most robust infiltration, possibly due to expression of the EGFRvIII neoantigen in the tumour cells (Figure 6C and Supplemental Figure 7). Importantly, immune infiltration was not due to tdTomato overexpression, as integration of a piggyBac construct encoding for tdTomato alone did not elicit an immune response (Supplemental Figure 6B, C). We therefore next assessed the immune microenvironment in bulk and margin by FACS and immunofluorescence analysis, again using EGFR tumours as a model. The overall proportion of Cd45 immune cells was significantly increased in the bulk, whereas the margin had levels of infiltration comparable to tumour-free brain tissue (Figure 6D and Supplemental Figure 6D and 7). This indicates that the bulk accounts for the majority of the tumour immune infiltrate whereas the margin may represent an immune-cold microenvironment. Differences in Cd45 cells were reflected in all immune components analysed, including microglia, macrophages, Cd4 T cells, Cd8 T cells, Tregs and natural killer cells, which were selectively enriched in the tumour bulk (Figure 6E and Supplemental Figure 6E and 7). This pattern was consistent with the hypothesis that increased immune activity in the bulk induces dormancy via interferon signalling. To test this more directly, we used two complementary approaches. First, we examined the spatial distribution of immune cells relative to H2B-GFP^+^ LRC by immunofluorescence. Although enriched in necrotic regions, microglia and macrophages were evenly distributed throughout the rest of the bulk (Supplemental Figure 6F). Instead, rare natural killer cells were restricted to necrotic regions (Supplemental Figure 6G). This suggests that neither cell type may be functionally linked to acquisition of LRC phenotypes, as the distribution of H2B-GFP^+^ cells was not uniform either within or outside necrotic patches (Figure 5B, Figure 6F and Supplemental Figure 6H). In contrast, Cd4 and Cd8 T cells formed clusters in multiple tumour areas, including around necrotic regions, which appeared to co-localise at least in part with H2B-GFP^+^ LRC-rich regions (Figure 6F and Supplemental Figure 6H).

To quantify a potential spatial correlation between these populations, we used digital pathology. We applied supervised and semi-supervised algorithms to identify the exact location of T cells and H2B-GFP^+^ LRC in immunofluorescence confocal tile scan images of the tumours (balanced accuracy: LRC = 0.96, T-cells = 0.95). The LRC population was subdivided according to GFP intensity as a surrogate for their proliferative status (Supplemental Figure 5D), with H2B-GFP^high^ cells being the least and H2B-GFP^low^ the most proliferative (unsupervised three-classes k means applied at each sample) and spatial relationships measured using cell-to-cell distance and abundance-based approaches (Figure 6G). Both Cd4 and Cd8 T cells were found to be closer to H2B-GFP^high^ than H2B-GFP^low^ cells (Figure 6H; GLMM, factor link type, T cell-GFP^high^ vs T Cell-GFP^low^: F = 193.467, p = 6.22e-44), while Cd8 T cells were closer to both H2B-GFP^+^ tumour populations than Cd4 T cells (Fig 6H; GLMM, factor T cell type Cd8 vs Cd4: F = 6.833, p = 0.039), in the absence of a significant interaction between these variables (GLMM, interaction between factor T cell type and link type: F = 0.045, p = 0.083). Furthermore, measurement of the Morisita-Horn overlap index revealed that the co-localisation of both Cd8 and Cd4 T cells with H2B-GFP^high^ was higher than 0 (Cd8: t[3] = 4.508, p = 0.02; ; Cd4: t[3] = 2.4, p = 0.047) within immune hotspots (defined by Getis-Ord’s G* on T cells distribution) and, controlling by the T cell/H2B-GFP^+^ ratio within immune hotspots, the co-localisation of H2B-GFP^high^ cells with Cd8 T cells was higher than with Cd4 T cells (Figure 6I, J; F[1,5] = 9.076, p = 0.029; 95% CI difference = 0.028 – 0.348). Outside of immune hotspots, only co-localisation of H2B-GFP^high^ cells and Cd8 T cells was significantly different than 0 (one-sided t-test t[3] = 4.048, p = 0.014; Cd4: t[3] = 1.73, p = 0.09), and no significant differences in co-localisation with H2B-GFP^high^ cells were detected between Cd4 and Cd8 T cells (Figure 6I, J; GLM F[1,5] = 1.39, p = 0.29). However, H2B- GFP^high^ tumour cells showed overall higher co-localisation with Cd4 or Cd8 T cells than H2B- GFP^low^ cells resampled to control for difference in abundance between the two GFP subpopulations (Supplemental Figure 6I). Together, this spatially-resolved quantification suggests that H2B-GFPhigh LRCs reside in close proximity to T cells, particularly, to the Cd8 T cell compartment.

Second, we functionally assessed the role of interferon signalling in driving tumour dormancy. EGFR-H2B-GFP tumours were induced in *Ifnar1^-/-^* or compound *Ifnar1^-/-^;Ifngr1^-/-^* mice, which are homozygous knock-out for type I or type I/II interferon signalling, respectively (Figure 6K) (Huang et al., 1993; Muller et al., 1994). Background-matched wildtype mice were used as controls. Following pulse-chase experiments, we measured the proportion of H2B-GFP^+^ LRC in the three cohorts and found a significant decrease in both mutant strains relative to control tumours, indicative of a reduction in dormant cells in the absence of interferon signalling (Figure 6L). Importantly, interferon signalling-deficient and wildtype tumours had comparable levels of immune infiltration, indicating that the phenotype was not due to reduced immune activity in the interferon mutant strains (Supplemental Figure 6J). Consistent with these findings, recombinant type I or II interferons, alone or in combination, were sufficient to decrease the proliferation of primary H2B-GFP^-^ tumour cells purified from EGFR tumours (generated in a wildtype background) *in vitro* (Figure 6M). Together, these results indicate that dormancy is at least in part a bulk phenotype induced by interferon signalling in T-cell-rich niches.

## Discussion

The GBM margin is notoriously difficult to study in patients due to the challenges of sampling and identifying invasive tumour cells, which, by definition, are left behind following surgical resection (Cuddapah et al., 2014; Vehlow and Cordes, 2013). To circumvent this problem, we developed three somatic mouse models of GBM driven by some of the most common combinations of the human mutations. Overall, the models show striking similarities with the human disease, recapitulating the histology, transcriptional and cellular heterogeneity, and immune microenvironment of patient tumours (Patel et al., 2014; Wang et al., 2017). We found that regardless of genetics, all tumours mirrored the main developmental lineages of the brain, containing cells of astrocytic, oligodendrocytic, and transit amplifying progenitor/neuronal progenitor fate, alongside an injured NPC-like state. These findings are remarkably consistent with recent findings by Neftel et al. and Couturier et al., in which human GBM cells were found to exist in astrocyte-like, oligodendrocyte-progenitor-like, neural progenitor-like and mesenchymal-like states (Couturier et al., 2020; Neftel et al., 2019). These similarities further underscore the accuracy of our models in recapitulating the human disease and support an NSC origin for GBM (Alcantara Llaguno et al., 2009; Lee et al., 2018).

Although driver mutations biased the frequency of specific fates within our models, as was observed in human tumours (Neftel et al., 2019), our results indicate that genetics play an overall modest role in driving tumour phenotypes. In contrast, we find that tumour region is a major determinant. Within the bulk, tumour cells of all genotypes progressed along an injury-like trajectory culminating with interferon-induced dormancy in T-cell niches. Intriguingly, this behaviour is reminiscent of the aged SVZ, where neural progenitors were shown to undergo quiescence in response to age-dependent inflammation of the niche, including the production of interferons by infiltrating T-cells (Dulken et al., 2019; Kalamakis et al., 2019). Interferons have also been previously linked to dormancy in a handful of cancer types, which in a murine B-cell lymphoma model was released by Cd8 T cells (Correia et al., 2021; Farrar et al., 1999; Liu et al., 2017; Liu et al., 2018). It is of note that both type I and type II interferons mediated dormancy in our models. Together with the observation that dormancy often co-localises with immune hotspots and necrotic regions, this points to a model whereby dormancy results from a combination of paracrine adaptive interferon *γ* signalling produced by infiltrating T cells and autocrine innate interferon *α*/*β* signalling triggered by activation of the STING pathway in response to T cell-mediated killing of neighbouring tumour cells (Zhu et al., 2019). It is tempting to speculate that dormancy may be a general tumour response to T cell infiltration induced by the inflammatory microenvironment of the tumour bulk and that interferon blockade could be a promising strategy for chemosensitising GBM.

At the margin, where the immune microenvironment resembled that of normal brain, cells followed a developmental-like tumour hierarchy biased towards astrocyte-like fate, even in Pdgfr tumours that display an intrinsic oligodendrocyte-like fate bias. Thus, although tumours are often compared to wounds that don’t heal, our findings suggest that injury programmes are mostly relevant to the bulk of the tumour and may not play a major role in driving margin phenotypes, with key implications for GBM treatment, including checkpoint blockade immunotherapy (Dvorak, 1986). In line with our findings, a recent study proposed that human GSCs exist in either a neurodevelopmental or an inflammatory state (Richards, 2021). It would be of great interest to explore whether there is a correlation between GSC state and their tumour region of origin in patients.

The dominance of the microenvironment in controlling tumour behaviour is further emphasised by our observation that within the invasive margin astrocyte differentiation is highly localised to specific brain regions. We recently reported that invasion into white matter promotes differentiation towards oligodendrocyte fate in patient-derived models (Brooks et al., 2021). The findings presented here corroborate the observation of increased lineage progression at the margin and further suggest that even within a specific brain region, differentiation trajectories are highly heterogeneous and dependent on local extrinsic cues. This is of clinical relevance as it suggests that location within the brain could be predictive of tumour biology and, potentially, response to treatment. Comprehensively defining invasive phenotypes in their spatial context may therefore identify key biological vulnerabilities for eradicating invasive cells and is an important direction for future studies.

Our work also provides two important insights into the biology of stem-like tumour cells. First, it suggests that GSCs resembling NSCs may not be slow-cycling or quiescent, as is the case for their normal counterparts. Indeed, in agreement with recent findings in patients, in our models, actively dividing NSC-like cells occupied the top of the tumour hierarchy (Couturier et al., 2020). This is perhaps not surprising given that normal quiescent NSCs are maintained by key tumour suppressors that become inactivated in GBM, including *p53* and *Cdkn2a* (Gil-Perotin et al., 2006; Meletis et al., 2006; Nishino et al., 2008). In further support of this idea, we also found that the majority of label-retaining tumour cells did not have NSC signatures, but rather represented more committed progenitors that were induced to exit the cell cycle by the injury-like microenvironment of the tumour bulk. This is indicative of key differences between developmental and tumour hierarchies and highlights the importance of injury/inflammatory response programmes in gliomagenesis (Brooks et al., 2021; Richards, 2021; Wang et al., 2017). Second, we found an equal distribution of aNSC-like cells between bulk and margin. While we cannot exclude the possibility that other subsets of stem-like tumour cells may be enriched at the invasive niche, this work suggests that invasive potential is a general property of most, if not all, tumour cell subpopulations. This is somewhat unexpected given that GSCs were shown to be more invasive than their non-stem counterparts (Cheng et al., 2011). It remains to be determined if astrocyte-like differentiation is beneficial for invasion or rather a by-stander effect driven by the surrounding normal brain microenvironment. Regardless, identifying biological vulnerabilities of the astrocyte-like state would be an important next step for the development of treatments that more effectively target the GBM margin residuum.

In summary, our work reveals fundamental differences between the tumour bulk and margin and suggests that analysis of the bulk is not directly informative of margin biology or treatment. It further suggests that combinatorial therapies that take these differences into account will be required to improve patient outcome in this devastating disease.

## Acknowledgements

This work was funded by Cancer Research UK (S.P., C.G.D., A.P., L.C.), the NIHR Biomedical Research Centre (M.C.) and the Fundació La Marató de TV3 (HH). We thank A. Berns for *Cdkn2a^-/-^* mice, M. Aguet for *Ifnar1^-/-^* and *Ifnar1^-/-^;Ifngr1^-/-^* mice, S. Pollard for constructs, B. Antolin-Fontes for cloning, G. Rodriguez-Esteban for reads mapping, M. Pathania for technical advice, A. Flanagan for histopathological advice, J. Manji for help with microscopy, Y. Guo, G. Morrow and B. Wilbourn for help with FACS and S. Marguerat for helpful discussion and critical reading of the manuscript.

## Author Contributions

Conceptualization, S.P.; Methodology, C.G.D., M.C., H.H., S.Q., Y.Y. and S.P.; Software, E.M., H.H., S.P.C., Y.Y.; Validation, C.G.D., A.P., L.C.; Formal analysis, C.G.D., E.M., A.P., G-C, S.P.C., L.C., F.R.; Investigation, C.G.D., M.C., A.P., F. G-C., L.C., S.R.; Resources, H.H., S.Q., Y.Y. and S.P.; Data Curation, E.M.; Writing - Original draft, C.G.D., M.C., E.M. and S.P.; Visualisation C.G.D., E.M., A.P., S.P.C. and S.P.; Supervision, H.H., S.Q., Y.Y. and S.P.; Project administration, S.P.; Funding acquisition, H.H., S.Q., Y.Y. and S.P.

## Declaration of Interests

HH is co-founder and shareholder of OmniScope. The rest of the authors declare no competing interests.

## Methods

### Animals

All procedures were performed in compliance with the Animal Scientific Procedures Act, 1986 and approved by with the UCL Animal Welfare and Ethical Review Body (AWERB) in accordance with the International guidelines of the Home Office (UK). *Trp53^fl/fl^* mice were obtained from the Jackson Laboratory (*Trp53^tm1Brn/J^*; Jax 008462) (Marino et al., 2000) and *Cdkn2a^fl^*^/*fl*^ mice were provided by A. Berns (*Cdkn2^atm2Brn/A^*) (Krimpenfort et al., 2001). *Trp53^fl/fl^* pups were used for modelling Nf1 and Pdgfr tumour models, *Cdkn2a^fl^*^/*fl*^ mice were used for the EGFR model. Wildtype C57Black/6 mice were purchased from Charles River and used for the EGFR H2B-GFP model. *Ifnar1^-/-^* and *Ifngr1^-/-^;Ifnar1^-/-^* mouse lines were provided by Michel Aguet (Huang et al., 1993; Muller et al., 1994). Female and male mice were used for tumour modelling. Mice were monitored daily and sacrificed when they began to show signs of disease and reached humane endpoints. To study the distribution of GSCs between bulk and margin, EdU (50mg/kg; Insight Biotech, sc-284628A) was injected 2 hours prior to sacrifice to label rapidly dividing cells in the brain. To identify dormant tumour cells in the EGFR H2B-GFP model, doxycycline (Sigma, D891) was administered through the drinking water (0.2% doxycycline:1% sucrose) immediately following plasmid injection. Doxycycline withdrawal was carried out for a minimum of 2 weeks to dilute the H2B-GFP reporter in actively cycling cells.

### *In vivo* Electroporation

Plasmids were injected into the ventricle of isoflurane-immobilized pups at postnatal day 2 using an Eppendorf Femtojet microinjector (Eppendorf, 5247000030), followed by electroporation (5 square pulses, 50 msec/pulse at 100V, with 850 msec intervals). PiggyBase (0.5ug/ul) and PiggyBac vectors at a molar ratio of 1:1 were diluted in saline (0.9% NaCl) containing 0.1% fast green (Sigma, F7258).

### Tissue preparation and immunohistochemistry

Animals were perfused (4% paraformaldehyde in PBS) under terminal anaesthesia, brains collected and post-fixed overnight at 4°C in PFA (4%). Vibratome sections (50μm) were prepared and stored in cryopreservative (glycerol:ethylene glycol:PBS 1:1:2) prior to immunohistochemistry. For staining, floating sections were permeabilised overnight (1% triton-X-100, 10% serum in PBS) at 4°C, incubated in primary antibody overnight (1% triton-X-100, 10% serum in PBS) at 4°C and for 3 hours in secondary antibody (0.5% triton-X-100, 10% serum in PBS) at room temperature. Sections were counterstained with DAPI (Insight Biotechnology, sc-3598) for 10 min at room temperature and mounted with antifade mounting solution (Prolong gold antifade mountant, Thermo Fisher, P36934). Images were acquired on Confocal LSM 880 (Zeiss) and analysed on ImageJ (RRID:SCR_003070).

The following antibodies were used: rabbit anti-Sox2 (1:500; Abcam, ab97959), rabbit anti-GFAP (1:1,000; Dako, Z0334), rabbit anti-Olig2 (1:500; Millipore, ab9610), rabbit anti-RFP (1:500; Antibodies Online, ABIN129578), goat anti-Sox9 (1:50; R&D AF3075), rat anti-Cd45 (1:500; BD, 550539), rat anti-Cd68 (1:500; Abcam, ab53444), rabbit anti-Iba1 (1:1,000; Wako, 019-19741), rat anti-Cd8a (1:250; Thermofisher, 14-0808-82), rat anti-Cd4 (1:100; BD, 550280), goat anti-Nkp46 (1:250; R&D, AF2225), rat anti-Cd31 (1:100; BD, 550274). For detection of EdU, sections were stained with Click-it EdU Alexa Fluor 647 Imaging Kit (Invitrogen, C10340) following manufacturer guidelines.

For histopathology assessment, brains were post-fixed in formalin overnight before tissue processing and paraffin embedding. 3μm sections were cut and stained with haematoxylin and eosin using standard methods.

### Derivation and culture of cell lines

Neural stem cells were isolated from *Trp53^fl/fl^* and *Cdkn2a^fl/fl^* pups as previously described (Ottone et al., 2014). Briefly, pup brains (P9-14) were collected and the lateral ventricles dissected out. Neural stem cells were isolated by enzymatic digestion using papain dissociation (Worthington, LK003178). Cells were seeded in NSC media (DMEM/F12 supplemented with N2 (1x), B27 lacking retinoic acid (1x), kanamycin (100mg/ml)/gentamycin (2mg/ml), heparin (4mg/ml), FGF (10ng/ul) and EGF (20ng/ul) and expanded as neurospheres for one passage prior to plating on laminin-coated (1:200 in PBS) plates and cultured in GSC media as previously described (Pollard et al., 2009).

For preparation of tumour cell lines from mice, brains from tumour-bearing animals were collected into ice-cold HBSS media. Under fluorescence guidance, tdTomato^+^ tumour regions were microdissected, enzymatically digested and cultured as described above, with the exception of EGFR tumour-derived cells which were maintained and subcultured as neurospheres throughout.

### Interferon treatment and cell proliferation assay

Proliferative GFP negative tumour cells were FACS-sorted from EGFR-H2B-GFP tumours after a 2-week doxycycline chase and cultured as neurospheres as above. Cells were incubated in the presence or absence of interferon *β* (1000 U/ml; R&D, 8234-MB-010), interferon *γ* (1000 U/ml; PeproTech 315-05-20) or both combined for 48 hours. EdU (10mM) was added to the media 2 hours prior to collection to label dividing cells. Following EdU labelling (10mM; Invitrogen, C10424) according to the manufacturer’s instructions, a minimum of 10,000 cells were analysed using a BD LSRFortessa X-20 Flow Cytometer and cell-cycle profiles measured using FlowJo software.

### Western Blot

GBM mutations were verified on tumour-derived and NSC cell lines by Western blotting. Protein lysates were prepared in RIPA buffer (containing protease (1:100; Sigma, P8340) and phosphatase inhibitors (1:500; Sigma, P5726 and P0044). Western Blots were performed following standard protocols. Membranes were incubated with primary antibodies in 5% milk in TBST (TBS+ 0.05% Tween) overnight at 4 °C, washed and incubated in secondary antibody (in 5% milk in TBST) at room temperature for 1h. Proteins were detected using Luminata Crescendo (Millipore, WBLUR0500) or Classico (Millipore, WBLUC0500) Western HRP reagents and imaged using the ImageQuant system.

The following primary antibodies were used: rabbit anti-p16 (1:500; Abcam, ab211542), rabbit anti-Trp53 (1:500; Novocastra Leica, NCL-L-p53-CM5P), goat anti-Pdgfra (1:500; R&D, AF1062), rabbit anti-Nf1 (1:1,000; Bethyl, A300-140A-M), rabbit anti-Pten (1:1,000, Cell Signalling, 9559), rabbit anti-EGFR (1:1,000; Millipore, 06847) and mouse anti-Gapdh (1:5,000; Abcam, ab8245). HRP secondary antibodies were purchased from ThermoFisher.

### Plasmid generation

Constructs were generated using InFusion Kit (Clontech, 638917) and T4 DNA Ligase (NEB, M0202S), following manufacturer’s guidelines. Plasmids were transformed in chemically competent bacteria strains Top10 (Thermo Fisher, C303003) and Stbl3 (Thermo Fisher, C737303). Stbl3 bacteria strain was used for PiggyBac vectors to minimise recombination. Plasmid construction and verification of constructs was designed using Snapgene software (RRID: SCR_015052). Previously described sgRNA were used to target *Pten* and *Nf1* (Zuckermann et al., 2015) for the Nf1 model and the *Cdkn2a* locus for the EGFR-H2B-GFP model (Weber et al., 2015).

### PiggyBases

The hGFAPMIN-PBase plasmid was generated by inserting the hGFAPMIN promoter from pAAV-GFAP-EGFP (a gift from Bryan Roth, Addgene # 50473) into pCAG-PBase plasmid (a gift from Paolo Salomoni) (Pathania et al., 2017). hGFAPMIN-SpCas9-T2A-PBase plasmid was generated by introduction of SpCas9-T2A into hGFAPMIN-PBase.

### PiggyBac plasmids

‘EF1α-tdTomato’ was a gift from Paolo Salomoni (Pathania et al., 2017). For the Nf1 model, hGFAPMIN and codon-improved Cre recombinase (iCre) sequences were inserted from hGFAPMIN-PBase and pBOB-CAG-iCre-SD (a gift from Inder Verma, Addgene # 12336) into EF1α-tdTomato PB vector. sgRNAs targeting Nf1 and Pten were cloned upstream of the EF1α- tdTomato sequence, as described above. For the Pdgfr model, hGFAPMIN-iCre sequence from Nf1 PB vector above and cloned into EF1α-TdTomato-CAG-Pdgfra (D842V) (a gift from Paolo Salomoni) (Pathania et al., 2017). For the EGFR model, EGFRvIII was PCR-amplified and cloned into the Pdgfr model PB vector to replace PdgfraD842V. To generate the EGFR-H2B- GFP PiggyBac plasmid, the EGFR model PB plasmid was modified as follows. hGFAPMIN- iCre sequence was replaced with a U6-sgRNA sequence targeting *Cdkn2a*. To introduce the H2B-GFP reporter, the tetracycline inducible expression construct was PCR-amplified from pCW57.1 (a gift from David Root, Addgene # 41393) and the H2B-GFP reporter was PCR-amplified from LV-GFP (a gift from Elaine Fuchs, Addgene # 25999) (Beronja et al., 2010) and cloned after the tdTomato sequence as a polycistronic construct with a T2A linker.

### Flow cytometry analysis

Brains were collected into ice-cold HBSS media and dissected into 1mm coronal sections using a brain matrix as above (WPI, RBMS-200C). The following regions were isolated under fluorescence guidance: tumour bulk, tumour margin and an equivalent area from the contralateral side (unless otherwise specified). Tissue was mechanically dissociated into small pieces, followed by enzymatic dissociation using Liberase TL (Roche, 05401119001) supplemented with DNAse (Sigma, 101041590001) for 30min at 37°C. Following addition of EDTA to stop the enzymatic reaction, cells were washed with PBS and filtered through a 70 µm cell strainer (Falcon, 352350) to remove large debris. After a blocking step in serum and Fc receptor blocking cocktail containing fetal bovine, mouse, rabbit and rat serums and anti- Cd16/32 antibody (BioXCell, BE0307) for 20 min on ice, cell suspensions were incubated with antibodies at 4°C for 20 min. For detection of intracellular epitopes, cells were fixed and permeabilised using BD CytoFix/CytoPerm kit (BD, 554714) for 20 min at 4°C in the dark for all panels, with the exception of the immune population panel where permeabilization and intracellular staining were performed for 2 h at 4°C in the dark. All centrifugation steps were carried out at 820g for 2 min and 820g for 5 min following permeabilization. Samples were acquired on a BD FACSymphony flow cytometer.

To compare the tumour populations from the bulk and the margin (Figure 4E), the following antibodies were used: rat anti-Cd45-BUV563 (Clone 30-F11, BD, 612924), rat anti-Cd11b- BUV661 (Clone M1/70, BD, 565080), mouse anti-β2-microglobulin-BUV805 (Clone S19.8, BD, 749215), mouse anti-MHC Class I H-2K^b^-BV510 (Clone AF6-88.5, Biolegend, 116523), rat anti-Bst2 (Cd317)-BV650 (Clone 927, BD, 747605), rat anti-MHC Class II (I-A/I-E)- BV711 (Clone M5/114.15.2, BD, 563414), rabbit anti-RFP (Antibodies Online, ABIN129578) with donkey anti-rabbit AF594, Viability dye-eFluor 780 (eBioscience, 65-0865-18). The following antibodies were used for analysis of the H2B-GFP LR population (Figure 5): rat anti- Cd74-BUV395 (Clone In-1, BD, 740274), rat anti-Cd45-BUV563 (Clone 30-F11, BD, 612924), rat anti-Cd11b-BUV661 (Clone M1/70, BD, 565080), rat anti-Ki67-eFluor450 (Clone SolA15, eBio, 48-5698-80), rat anti-Bst2-BV650 (Clone 927, Biolegend, 127019), rat anti-Cd44-BV786 (Clone IM7, Biolegend, 103059), rat anti-GFP-AF488 (Clone FM264G, Biolegend, 33807), rabbit anti-RFP (Antibodies Online, ABIN129578) with donkey anti-rabbit AF594, Viability dye-eFluor 780 (eBioscience, 65-0865-18). For analysis of the immune microenvironment, the following antibodies were used: mouse anti-Nk1.1-BUV395 (Clone PK136, BD, 564144), rat anti-Cd4-BUV496 (Clone GK1.5, BD, 564667), rat anti-Cd45- BUV563 (Clone 30-F11, BD, 612924), rat anti-Cd11b-BUV661 (Clone M1/70, BD, 565080), rat anti-Cd3-BUV737 (Clone 17A2, BD, 564380), rat anti-Cd8a-BUV805 (Clone 53-6.7, BD, 564920), rat anti-FoxP3-eFluor 450 (Clone FJK-16S, eBioscience, 48-5773-82), Viability dye-eFluor 780 (eBioscience, 65-0865-18).

Data was analysed using Flowjo (v10.7.1; RRID:SCR_008520). Data was compensated and only viable singlets were used for downstream analysis. For the analysis of tumour cells, Cd45 and Cd11b markers were used to exclude the hematopoietic compartment. Non-hematopoietic cells were gated based on tdTomato expression and a minimum of 1,500 cells (tdTomato^+^ cells for bulk and margin regions, and tdTomato^-^ for contralateral region) were used for further analysis. UMAP visualization of GFP, Cd44 and Bst2 markers was performed on concatenated data from 5 tumours (McInnes et al., 2018). Positive populations were manually gated based on fluorescence minus one controls (FMO) and projected onto UMAP to compare marker distribution between GFP^+^ and GFP^-^ populations.

To study the immune cell infiltration, Cd45 and Cd11b were used to identify the hematopoietic compartment. Main immune populations were manually gated as follows: Macrophages (Cd45^high^ Cd11b^+^), Microglia (Cd45^low^ Cd11b^+^), Cd8 T cells (Cd45^+^ Cd11b^-^ Cd3^+^ NK1.1^-^ Cd8^+^), Cd4 T cells (Cd45^+^ Cd11b^-^ Cd3^+^ NK1.1^-^ Cd4^+^), Cd4 Teff (Cd45^+^ Cd11b^-^ Cd3^+^ NK1.1^-^ Cd4^+^ FoxP3^-^), Cd4 Tregs (Cd45^+^ Cd11b^-^ Cd3^+^ NK1.1^-^ Cd4^+^ FoxP3^+^), Natural killers (Cd45^+^ Cd3^-^ Nk1.1^+^), Natural killer T cells (NKT) (Cd45^+^ Cd3^+^ Nk1.1^+^). See Supplemental Figure 7.

### Fluorescence-activated cell sorting for collection of single cells for RNA-sequencing

To collect single tumour cells for scRNA-seq, brains containing tumours were collected and dissected into 1 mm coronal sections using a brain matrix (WPI, RBMS-200C). The tumour bulk and invasive tumour front migrating into the striatum (margin) were micro dissected from the sections under fluorescence guidance. Brain regions were enzymatically dissociated to single cells using papain dissociation, as described above. Cells were resuspended into FACs buffer supplemented with RNAse inhibitors (2.5 mM HEPES, 1 mM EDTA, 1.5% BSA, 2.5% RNAse) and DAPI was added 5 min prior sorting (1:10,000; Insight Biotechnology, sc-3598). Fluorescence-activated cell sorting was performed on a BD FACSAria Fusion Class II Type A2 Biosafety Cabinet. Control tissue was processed in parallel to determine gating for the tdTomato^+^ tumour cells. These cells were sorted into 96-well plates containing RNA lysis buffer. For quality control purposes, half of each plate was sorted with tumour cells from the margin and the other half with tumour cells from the bulk, leaving one empty well. After sorting, plates were snap frozen on dry ice and then stored briefly at -80°C until library preparation.

### Single cell RNA library preparation

Full-length single-cell RNA-seq libraries were prepared using the Smart-seq2 protocol with minor modifications (Picelli et al., 2014). Briefly, freshly harvested single cells were sorted into 96-well plates containing the lysis buffer (0.2% triton-X-100, 1U/µl RNase inhibitor (Takara Bio, 2313A). Reverse transcription was performed using SuperScript II (ThermoFisher Scientific, 18064014) in the presence of 1 μM oligo-dT30VN (IDT), 1 μM template-switching oligonucleotides (QIAGEN), and 1 M betaine (Sigma 61962). cDNA was amplified using the KAPA Hifi Hotstart ReadyMix (Kapa Biosystems KK2601) and IS PCR primer (IDT), with 24 cycles of amplification. Following purification with Agencourt Ampure XP beads (Beckmann Coulter, A63881), product size distribution and quantity were assessed on a Bioanalyzer using a High Sensitivity DNA Kit (Agilent Technologies 5067-4628). A total of 140 pg of the amplified cDNA was fragmented using Nextera XT DNA sample preparation kit (Illumina FC-131-1096) and amplified with Nextera XT indexes (Illumina FC-131-1001). Products of each well of the 96-well plate were pooled and purified twice with Agencourt Ampure XP beads. Final libraries were quantified and checked for fragment size distribution using a Bioanalyzer High Sensitivity DNA Kit. Pooled sequencing of Nextera libraries was carried out using a HiSeq2500 (Illumina, RRID:SCR_016383) to an average sequencing depth of 0.5 million reads per cell. Sequencing was carried out as paired-end (PE75) reads with library indexes corresponding to cell barcodes.

### Single cell RNA-seq data analysis

#### Data pre-processing

After sequencing, libraries were inspected with the FastQC suite to assess the quality of the reads. Reads were then demultiplexed according to the cell barcodes and mapped on the mouse reference genome (Gencode release 21, GRCm38 (mm10)) with the RNA pipeline of the GEMTools 1.7.0 suite using default parameters (6% of mismatches, minimum of 80% matched bases, and minimum quality threshold of > 26) (Marco-Sola et al., 2012). For all samples, cells with <65% of mapped reads or <100,000 of total mapped reads were discarded. Cells in the 95% percentile of the distribution of detected genes were included in the downstream analysis, resulting in read count matrices containing 957 (EGFR), 1033 (Pdgfr) and 834 (Nf1) cells. Genes that were expressed in fewer than five cells were removed.

#### Clustering

Filtering, normalization, selection of highly variable genes (HVG), clustering and genotype integration of cells were performed according to the Seurat package (version 2.3.4) (Butler et al., 2018). Through this pipeline, read counts were log-normalized for each cell using the natural logarithm of 1 + counts per ten thousand. To avoid spurious correlations, genes were scaled and centered after library sizes were regressed out. These scaled z-scores values are then used as normalized gene measurement input for the clustering and to visualize differences in expression between cell clusters. Selection of HVG was based on the evaluation of the relationship between gene dispersion and the log mean expression (with default parameters), while their total number was limited to 3000 genes, which was close to the average of genes per cell in EGFR and Nf1 models, while Pdgfr cells displayed around 5000 genes.

The clustering procedure projects HVG onto a reduced dimensional space before grouping cells into subpopulations by computing a shared-nearest-neighbours (SNN) based on Euclidean distance. The clustering algorithm is a variant of the Louvain method, which uses a resolution parameter to determine the number of clusters (Waltman and van Eck, 2013). The resolution parameter was set depending on both the observed heterogeneity and the biological interpretation of the resulting clusters. At this step, the dimension of the subspace is represented by the number of significant principal components (PC), which was decided based on the distribution of the PC standard deviations and by the inspection of the ElbowPlot graph. Cluster identities were assigned using previously described genes and cluster-specific markers obtained by differential expression analysis. UMAPs were used to visualize clusters and gene expression of biological relevant markers and signatures.

#### Data integration

After the three GBM models were analysed and annotated independently, we integrated them to find common patterns between them. The integration was performed by using the Seurat package, by which is possible to identify biological corresponding cells (anchors) between pairs of data sets, allowing data harmonization and comparison across tumours of different genotype. The algorithm makes use of the Canonical Correlation Analysis (CCA), a method that is able to learn gene correlation structures that are conserved across datasets (Hardoon et al., 2004). To do that, it identifies a fixed number of genes (i.e. the anchor feature parameter; in this case we used 6000 genes) that are then used to find relationships between cells across the different data sets.

#### Differential Expression and GO Analysis

Cluster-specific markers were identified through the Seurat function FindAllMarkers using the Wilcoxon’s rank sum test. The top 100 positive markers of each cell type were used as the signature for that type in order to compare them with external signatures. To visualize the similarity between cell type annotations from other studies, we applied matchSCore2 (Mereu et al., 2020), which computes Jaccard Index to quantify the overlap between cell-type signatures. Gene Ontology enrichment analysis was performed with the simpleGO package.

#### Trajectory analysis

Trajectory analysis and pseudo-ordering of cells was performed with the Monocle (Qiu et al., 2017) package (version 2.8.0) using the previously identified HVG from each individual analysis as ordering genes. Gene counts were modelled using the negative binomial distribution (negbinomial) as defined in the family function from the VGAM package. As for the clustering, the expression space was reduced down before ordering cells using the “DDRTree” algorithm, which allows 2D visualization and interpretation of the trajectory of cell states transitions through the provided set of genes.

#### Alignment of single-cell trajectories

To compare the single-cell expression dynamics observed in the trajectory analysis by each individual model, we have applied the cellAlign package (Alpert et al., 2018). CellAlign enables the alignment of two pseudotime ordering by a quantitative framework that relies on time warping algorithms. In doing that the tool assumes that starting and ending points of the trajectories are matching, as it was observed in the integrated trajectory analysis of the three GBM models. Briefly, individual pseudotime values assigned to each cell are divided into equally distributed points (meta-cells) along the trajectory to avoid data sparsity associated with single-cell data. Gene expression of meta-cells is averaged and their Euclidean distances are used to identify matches between trajectories. The resulting distance matrix is then used to represent the similarity between two trajectories. A line that minimizes the overall alignment-based distance is displayed to recapitulate the changes along the trajectory. Identical trajectories for example would match in each meta-cell and thus the resulting alignment-based distance will be zero. In this case the line would be diagonal and go from the starting point of the trajectory in the upper left of the distance matrix to the end point in the lower right. Any deviation represented in this line indicates a shift in the pseudotime resulting from comparative alignment. We used EGFR as a reference data set for the pairwise comparison with the other two models.

### Digital pathology

#### Cell detection and validation

We applied supervised and semi-supervised algorithms to identify the exact location of T cells and H2B-GFP^+^ LRC in immunofluorescence confocal tile scan images of the tumours. From each confocal image, we extracted the channels of GFP (H2B-GFP LRC), AF647 (T cells), and DAPI. All analyses were carried out using QuPath (Bankhead et al., 2017) and ImageJ (Schneider et al., 2012) software. For T cells, we run a cell segmentation and trained a supervised Random Trees classifier with 1140 annotations for training and 742 annotations for validation from non-overlapping regions with the training annotations made by CGD and LC. To detect GFP tumour cells, we implemented a Random Trees classifier with a semi-supervised pipeline, allowing to increase the detectability while maintaining the original label intensity of tumour cells. For the semi-supervised algorithm, we first trained a Random Trees classifier (classifier gfp v.1) with 1000 annotations on GFP cancer cells and 1000 annotations for the background (made by CGD and LC). As the classifier gfp v.1 includes the bias of the observer, it is not able to detect low-intensity GFP cells; then on the same tile, we increased the brightness and contrast (automatic B&C ImageJ). We applied the classifier gfp v.1 on that image and saved the predictions that served as new annotations for the original GFP (non-auto B&C) allowing us to train a new classifier (classifier gfp v.2). That approach maximises the detection of cells with lower intensities (fast-cycling cells). To validate the classifier gfp v.2, CGD and LC made 860 independent annotations in non-overlapping regions with the training annotations. To compute the balanced accuracy of the T and GFP cell classifiers, we obtained a binary mask for the predicted cells by each algorithm. We quantified true positive, true negative, false positive or false negative frequencies according to the value of the binary mask for the annotations’ coordinates (caret R package) (Kuhn, 2008). Finally, we run a simple cell segmentation on the DAPI channel and obtained a dilated binary mask that allowed us to remove detected GFP cells without DAPI marker.

This detection method allowed us to obtain the location of each cell within the sample. Only for GFP tumour cells, we saved features related to intensity as a surrogate of the proliferative status at single-cell resolution to assess spatial relationships of dormant and proliferative tumour cells with immune cells. Due to the similarity between Cd4 and Cd8 markers, we use the same algorithm for immune cells detection. All the images were formatted to 8-bit with intensity values ranging from 0-255. To reduce biases, CGD and LC were not directly involved in the implementation of this pipeline beyond their annotations.

#### Spatial metrics for co-localisation of T cells and H2B-GFP LRC

We identified LRC and proliferative phenotypes by applying unsupervised k-means clustering with k=3 on the single-cell maximum intensity value. This identified GFP cell with low, medium and high intensity. The clustering was applied independently to each sample. This allowed us to examine the spatial relationship between T cells and GFPhigh and GFPlow cells through a distance-based approach and an abundance-based approach (Maley et al., 2015).

The distance-based approach consists of representing the distribution of H2B-GFP+ cells (GFPhigh and GFPlow) and T cells in each sample as a network, where cells are the node and the distance between neighbouring cells are the links. For each sample, we run a Delaunay triangulation algorithm allowing us to obtain the spatial network and the distance between cells. We evaluated if the distance between cells differed between the two classes of links that connect (1) T cells and GFPlow cells and (2) T cells and GFP-high cells. As a second explanatory variable, we compared the distance between (1) Cd4^+^ and GFP cells and (2) Cd8+ and GFP cells, grouping GFPlow and GFPhigh cells. With these two explanatory variables, link class and T cells, we built a linear mixed model (lme4 R package) (Bates et al., 2015), with the logarithm (log10) of the distance as the response variable, link class and T cells as fixed factors, link class nested in T cells, and the sample as an explanatory variable with a random effect. If the null hypothesis for the fixed factor is rejected, we evaluate *a posteriori* comparisons between the corresponding factor levels applying the Satterthwaite method for the computation of residual degrees of freedom.

For the abundance-based approach we computed a discrete colocalisation measure based on the application of Morisita’s dispersion and Morisita-Horn overlap indices^4^ on the local co-occurrence of Cd4 T cells and GFPlow or GFPhigh cells. For each sample, we computed the Morisita dispersion index (Eq 1) at different spatial scales defined by the number of square quadrants or patches implemented by the R package IDmining (Golay et al., 2014), that measures the degree of randomness in cell distribution.

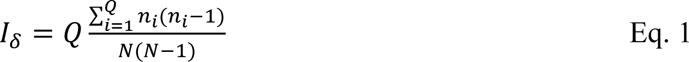

The algorithm subdivides the region of interests in *Q* quadrants or patches with a value of the diagonal (δ) and computes *I*_δ_ based on the abundance of cells (*n*_*i*_) in the patch and the total number of cells or points (N). We iterated the algorithm from one to 90K subdivisions (patches) for each sample and took the value of the diagonal that maximises *I*_δ_, as the distance where the spatial pattern diverges the most from complete spatial randomness. The value, which is sample-dependent, was used to create a polygonal grid for each sample and compute the Morisita-Horn overlap index (Eq 2) that calculates the probability to detect two classes of cells, for simplicity x and y, in the same quadrant with a similar relative abundance

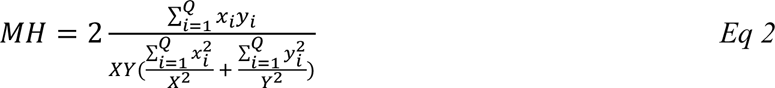

Where xi and yi are the quadrant abundances of the classes and X and Y are the sample abundance of the classes. We computed the MH index at two scales within T cell hotspots identified computing Getis-Ord G* and outside these hotspots. We tested with a one-sided t-student if the observed MH GFPhigh-Cd4 or GFPhigh-Cd8 differs from 0 (null hypothesis is that the observed colocalisation matches the expected for a random distribution). Additionally, with a general linear model (GLM), we tested if the MH GFPhigh differs between T cell types, adding the T cells/GFP ratio as a covariate. The statistical evaluation was made independently at both scales (inside and outside of immune hotspots). The normality of the variables, raw and residuals, was confirmed with a Shapiro-Wilk normality test.

#### Assessment of colocalisation between T cells and GFP subpopulations

Within samples, the abundance of GFPlow and GFPhigh cells is expected to differ because a relative minority of tumour cells remains low-cycling (GFPhigh). To compute comparable MH indices between T cells and GFP subpopulations (low and high) we therefore controlled for differences in abundance to rule out density biases. After patch detection with the Morisita dispersion index (Eq 1), we computed the Morisita-Horn overlap index for randomly sampled GFPlow cells where their abundance equals the observed abundance of GFPhigh. For each sample, random sampling was run 500 times; hence obtaining 500 values of MH between GFPlow and the corresponding T cell class. We compute a z-test to evaluate if the observed MH GFPhigh-Tcell is higher than the mean MH GFPlow-Tcell index from the random resampling for each sample.

### Data availability

scRNAseq data generated for this study has been deposited to GEO and will be made publicly available following publication.

### Statistical analysis

All statistical analysis were performed using Prism (GraphPad, RRID:SCR_002798). Mantel-Cox log-rank test was performed for survival data statistical analysis. Statistical tests and significance are described in figure legends (p values = * <0.05, ** <0.01, *** < 0.001 ****<0.0001). Shapiro-Wilk normality test was used to test normal distribution of samples. If no statistical significance is indicated on a graph, then ns > 0.05.

