## Supplemental Figures for "Glioblastoma cell fate is differentially regulated by the microenvironments of the tumour bulk and infiltrative margin"

### Supplemental Figure 1

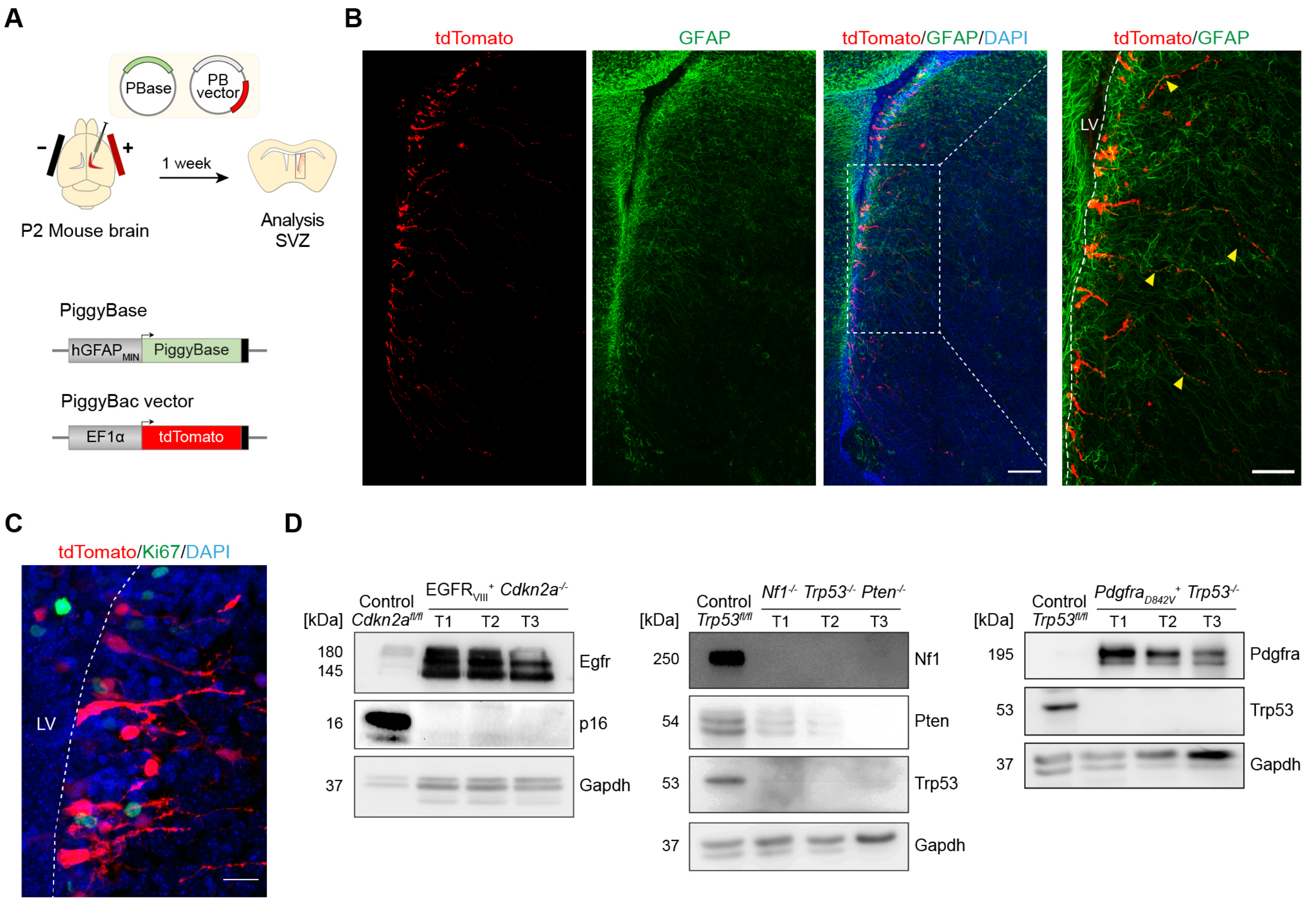

**Supplemental Figure 1. hGFAP reporter specificity and mutation targeting. A**, Schematic of experimental set up. **B** and **C**, representative immunofluorescence staining for GFAP (B) and Ki67 (C) of the lateral ventricles of pups 1 week following electroporation with a tdTomato reporter targeted to NSCs by the $hGAFP_{MIN}$ promoter. Note the characteristic radial glia morphology of targeted cells (arrowheads). Image on the right in B is the magnification of the boxed area on the left. Scale bars=200µm, 100µm for B and 20µm for C. **D**, Western analysis of levels of the indicated proteins in normal neural stem cells (NSC) and primary tumour cells isolated from three independent EGFR (left), Nf1 (middle) and Pdgfr (right) models. Related to Figure 1.

### Supplemental Figure 2

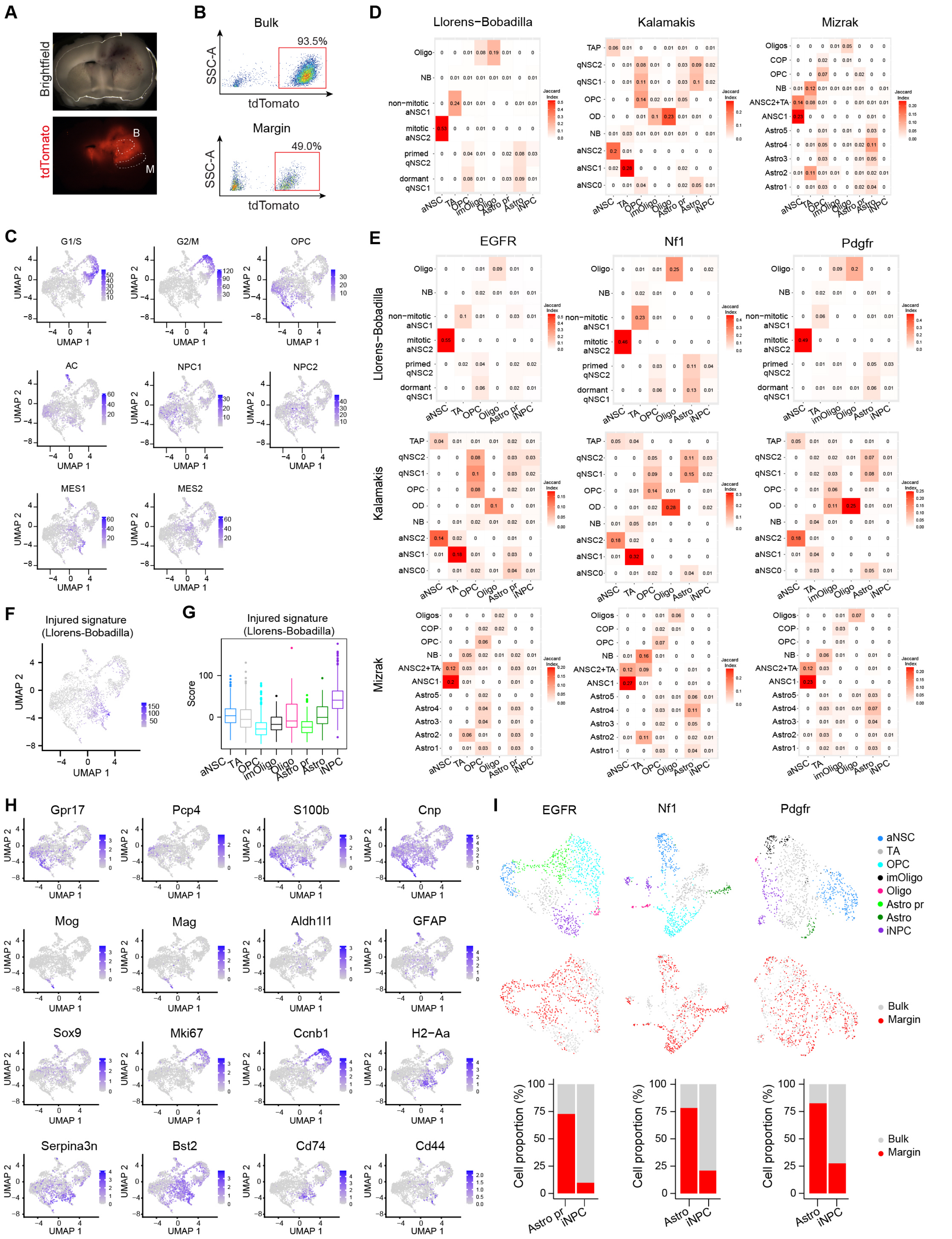

**Supplemental Figure 2. Derivation of tumour cell signatures.** **A**, Brightfield and epifluorescence images of a representative Nf1 tumour indicating bulk (b) and margin (m) regions of microdissection. **B**, FACS analysis of tdTomato$^+$ tumour cells in the bulk and margin of the tumour shown in A. Note the lower proportion of tumour cells in the margin. **C**, UMAP visualisation of the 2824 cells from the combined GBM tumour models. Signatures from Neftel et al. are highlighted in blue according to normalised expression levels, as follows. Proliferative cells (G1-S and G2-M), Oligodendrocyte progenitors (OPC), Astrocyte (AC), Neuronal progenitors (NPC1 and NPC2), Mesenchymal subtypes (MES1 and MES2). **D**, Jaccard Indexes of cell type specific markers identified in the integrated dataset and cell population signatures from Llorens-Bobadilla et al., Kalamakis et al and Mizrak et al. For each annotated cell population, the top 100 ranked markers were used. **E**, Jaccard Indexes of cell type specific markers defined for each individual tumour model separately and compared with cell population signatures from the Llorens-Bobadilla, Kalamakis and Mizrak studies. For each annotated cell population, the top 100 ranked markers were used. **F**, UMAP visualisation of the 2824 cells from the combined GBM tumour models. Injured NPC signature from Llorens-Bobadilla et al is highlighted in blue. Colour intensity represents the sum of z-scores among the genes in the signature. **G**, Boxplots comparing injured signature scores across cell populations in the combined tumour dataset. Boxplots display the minimum, 1st, 2nd, 3rd quartile and maximum of signature scores obtained for the different cell types. **H**, UMAP visualisation of normalized expression levels for selected marker genes of the distinct tumour subpopulations in the combined dataset. Shown are markers for OPCs (Gpr17, Pcp4), Oligodendrocyte progenitors (S100b, Cnp), Oligodendrocytes (Mog, Mag), Astrocytes (Aldh1l1, Gfap), Astrocyte progenitors (Sox9), aNSC (Mki67, Ccnb1), iNPC (H2-Aa, Serpina3n, Bst2, Cd74, Cd44). **I**, UMAP visualization of the clustering analysis of the separate tumour models. Cells are coloured by cell type (top) and tumor region (below). Cell proportions of bulk and margin regions are shown (bottom) for Astro pr and iNPC subpopulations. Related to Figure 2.

### Supplemental Figure 3

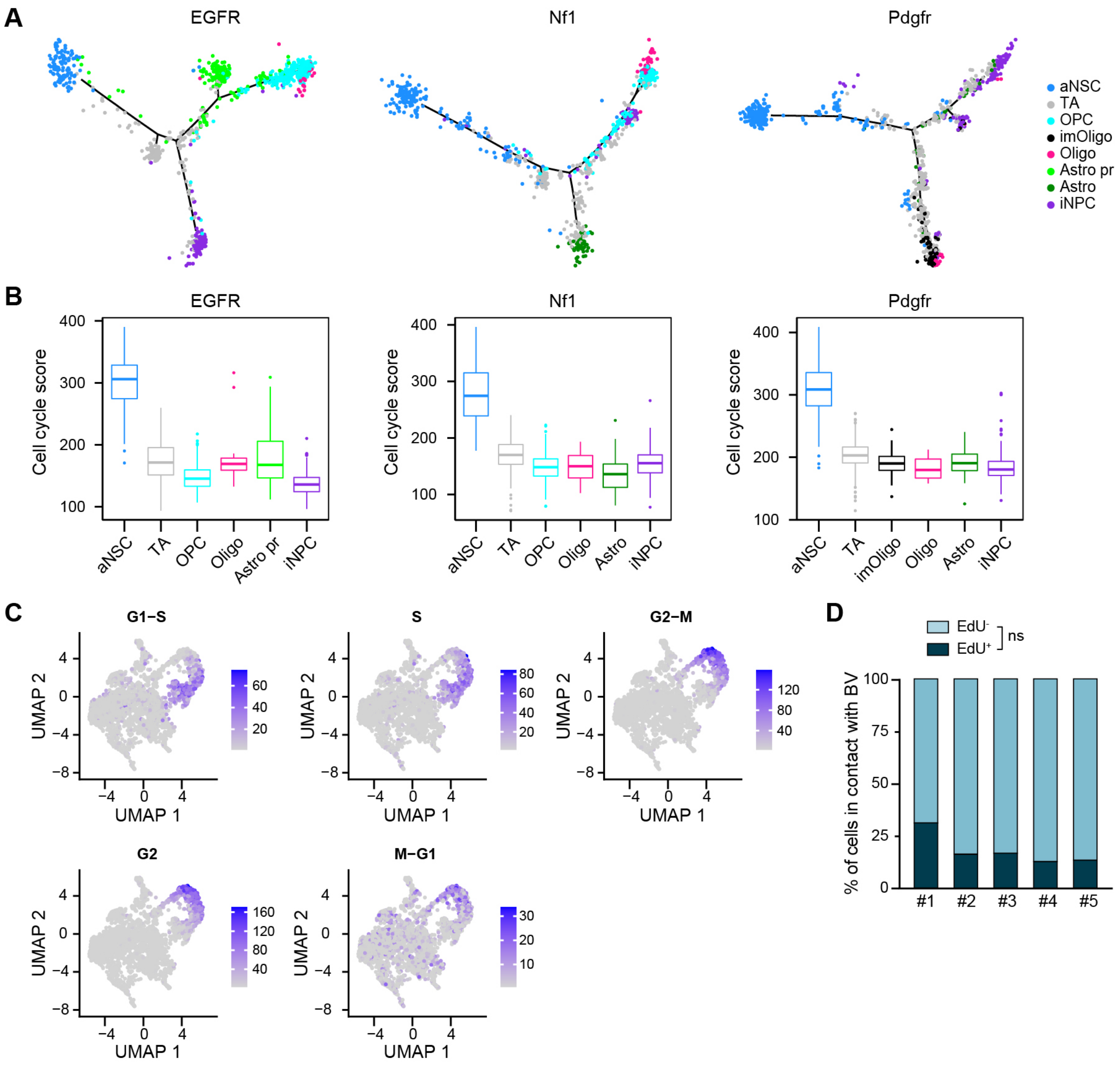

**Supplemental Figure 3. Proliferative aNSC-like cells fuel the tumour hierarchy. A**, Differentiation trajectory analysis of individual GBM models. Cells are coloured by cell type. Cell types were defined using clustering analysis of each model separately. **B**, Boxplots comparing the cell-cycle signature score across tumour cell populations in the three separate tumour models. Boxplots are ordered from higher to lower cell cycle score and display the minimum, 1st, 2nd, 3rd quartile and maximum of scores for each cell type. **C**, UMAP visualisation of the 2824 cells from the combined GBM tumour models. Cell cycle phase signatures are highlighted in blue. Colour intensity represents the sum of z-scores among the genes in the signature. **D**, Quantification of the percentage of $EdU^+$ and $EdU^-$ cells associated with blood vessels at the tumour margin of EGFR tumours. n=5, Three ROIs per tumour were counted. Each independent repeat is shown. Wilcoxon matched-pairs signed rank test (ns: p=0.0625). Related to Figure 3.

### Supplemental Figure 4

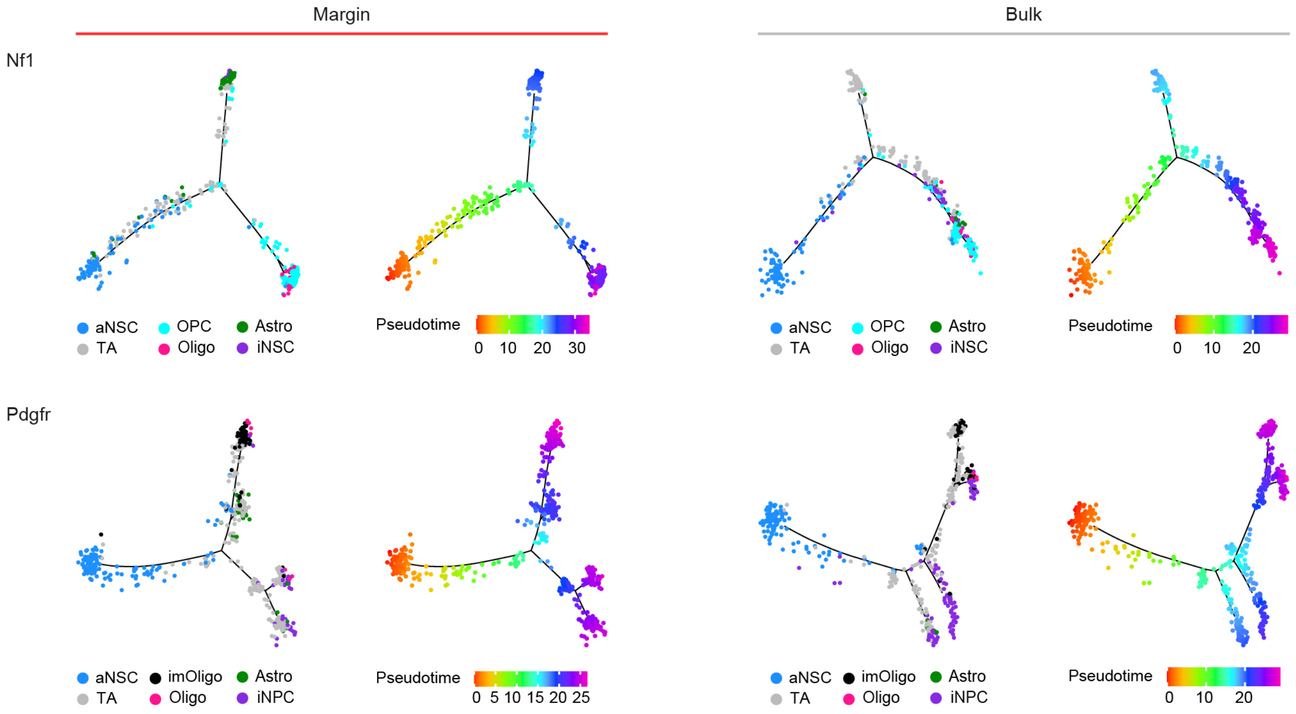

**Supplemental Figure 4. Fate choice differs between bulk and margin.** Differentiation trajectory analysis of 463 margin (left) and 494 bulk (right) cells in Nf1 tumours (top) and of 463 margin (left) and 494 bulk (right) cell in Pdgfr tumours (bottom) coloured by cell type. Related to Figure 4.

#### Supplemental Figure 5

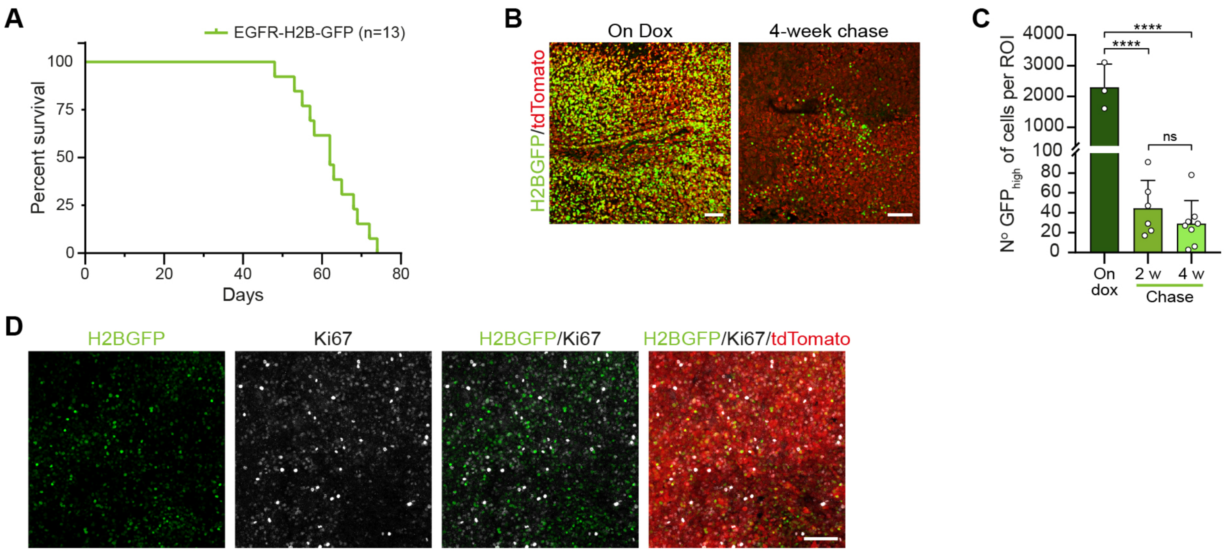

**Supplemental Figure 5. Validation of EGFR-H2B-GFP model.** **A**, Kaplan-Meier survival plots for the EGFR H2B-GFP model. Median survival of 62 days. n=13. **B**, Representative fluorescence images of EGFR-H2B-GFP tumours immediately after a 5 week doxycycline pulse (on Dox) and 4 weeks after chase. Scale bars=100 µm. **C**, Quantification of the number of H2B-GFP$^{+}$ tumour cells in EGFR-H2B-GFP tumours immediately after a 5-week doxycycline pulse (on Dox) and after a 2- (2w) and 4-week (4w) chase. n=6 ROIs from 2 tumours for 2w and 8 ROIs from 3 tumours for 4w. One-way ANOVA with Tukey's multiple comparison test (****: $p < 0.0001$, ns: $p=0.9945$). Mean±SD. **D**, Representative fluorescence images of EGFR-H2B-GFP tumours chased for two weeks and stained for Ki67. Note how all H2B-GFP$^{high}$ and most of H2B-GFP$^{low}$ cells are negative for Ki67. Scale bar=100 µm. Related to Figure 5.

### Supplemental Figure 6

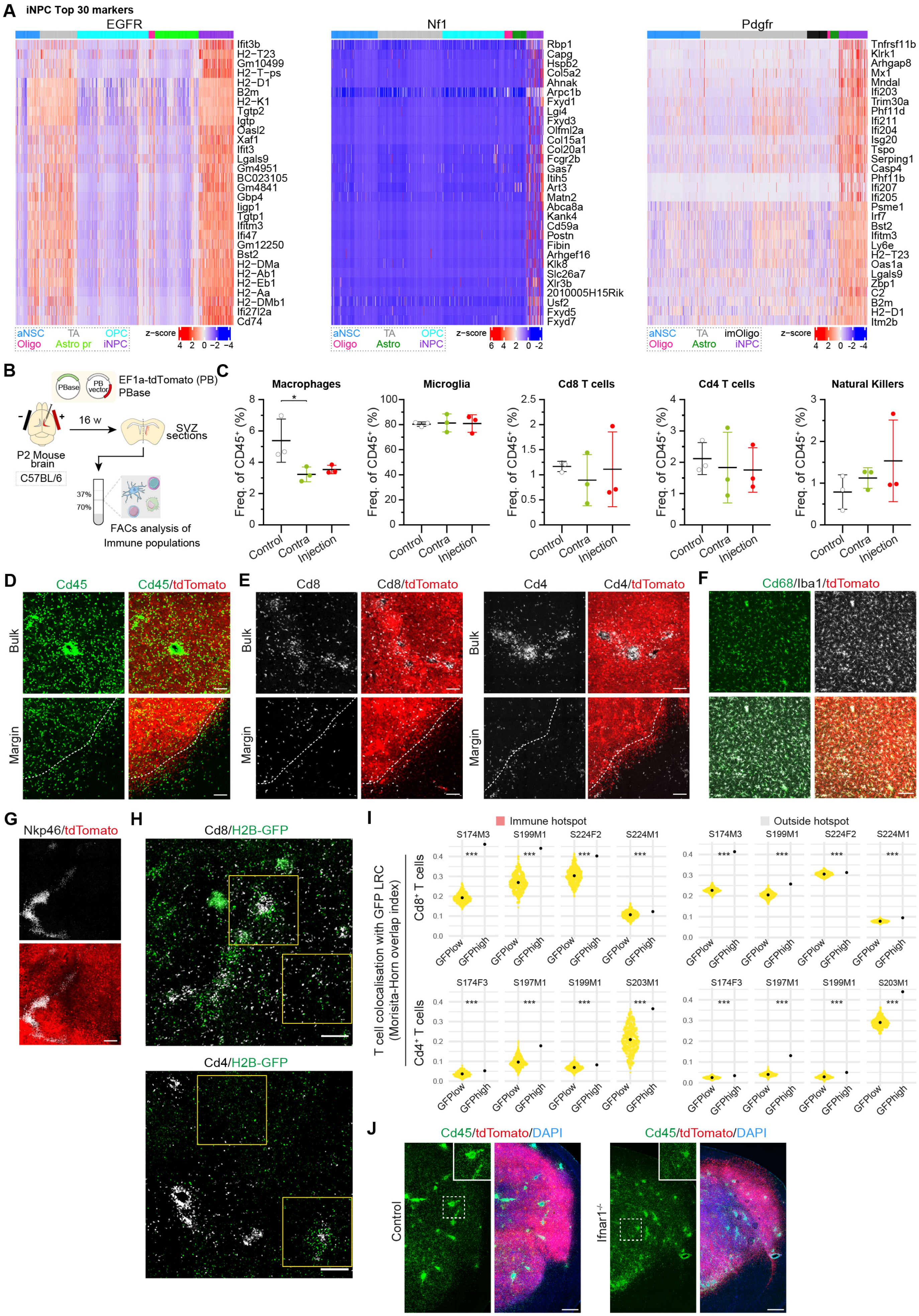

**Supplemental Figure 6. Characterisation of the immune microenvironment of the tumour models.** **A**, Heatmap of the top 30 markers of the iNPC population in each GBM model. Columns are grouped by cell type (top bar). Normalized gene expressed values (z-scores) are shown. **B**, Schematic of experimental set up to assess tdTomato immunogenicity. The injected and contralateral hemisphere were separated under fluorescence guidance and subjected to a 37-70% Percoll gradient to enrich the immune cell fraction. **C**, FACS quantification of the indicated main immune populations in the electroporated (Injection) and contralateral (Contra) hemisphere and intact control brains (Control) relative to the hematopoietic compartment ($Cd45^{+}$). n=3 brains per group. Mean ± SD. One-way ANOVA with Tukey's multiple comparison test for Macrophages (Control vs Contra: p=0.0494, Contra vs Injection: p=0.8991, Control vs Injection: p=0.0860), Microglia (Control vs Contra: p=0.9793, Contra vs Injection: p=0.9955, Control vs Injection: p=0.9940), Cd8 T cells (Control vs Contra: 0.8056, Contra vs Injection: p=0.8714, Control vs Injection: p=0.9904), Cd4 T cells (Control vs Contra: p=0.9064, Contra vs Injection: p=0.9929, Control vs Injection: p=0.8553). Kruskal-Wallis with Dunn's multiple comparison test for NK cells (Control vs Contra: p=0.6933, Contra vs Injection: p>0.9999, Control vs Injection: p=0.6097). **D**, Representative immunofluorescence staining for Cd45 (green) of the bulk and margin of EGFR tumours. Scale bar=100µm. **E**, Representative immunofluorescence staining for Cd8 (grey, left) and Cd4 (grey, right) of the bulk and margin of $tdTomato^{+}$ (red) EGFR tumours. Scale bars=100 µm. **F**, Representative immunofluorescence staining of the bulk of an EGFR tumour for the microglia and macrophage markers Iba1 (grey) and Cd68 (green). Scale bar=100µm. **G**, Representative staining for the NK cell marker Nkp46 of the bulk of an EGFR tumour. Scale bar=200µm. **H**, Representative Cd8 (top) and Cd4 (bottom) immunofluorescence staining of the EGFR tumour bulk. Scale bar= 200µm. Yellow boxes denote regions that are magnified in Figure 6F. **I**, Measurement of Morisita-Horn overlap indexes of $H2B\text{-}GFP^{high}$ (GFPhigh) or $H2B\text{-}GFP^{low}$

(GFPlow) tumour cells and Cd8 (top) and Cd4 (bottom) T cells. To balance abundance between H2B-GFP$^{low}$ and H2B-GFP$^{high}$ phenotypes, the H2B-GFP$^{low}$ population was randomly resampled 500 times, the central dot for GFP$^{low}$ indicates the mean value. n=500 null replicates for T cells-H2B-GFP$^{low}$ colocalization. z-test. **J**, Representative immunofluorescence staining for Cd45 of EGFR tumours generated in *Ifnar1*$^{-/-}$ and isogenic wildtype control. Scale bar=500 μm. Related to Figure 6.

### Supplemental Figure 7

**A**

**Gating strategy to study tumour populations**

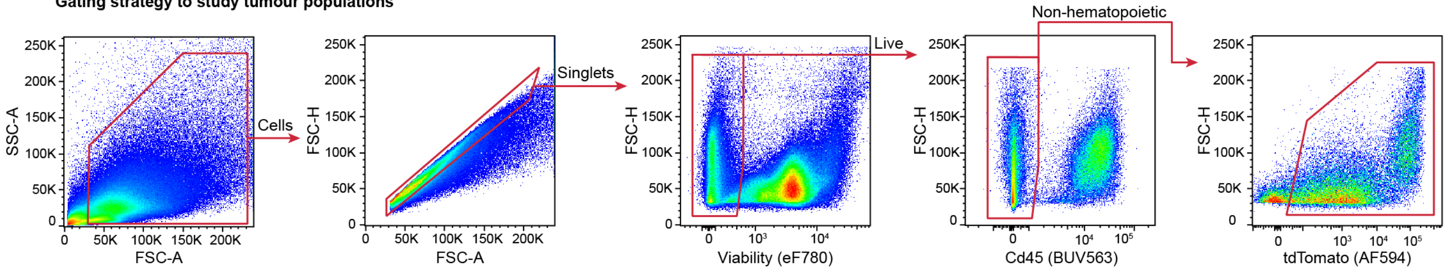


**B**

**Gating strategy to study immune populations**

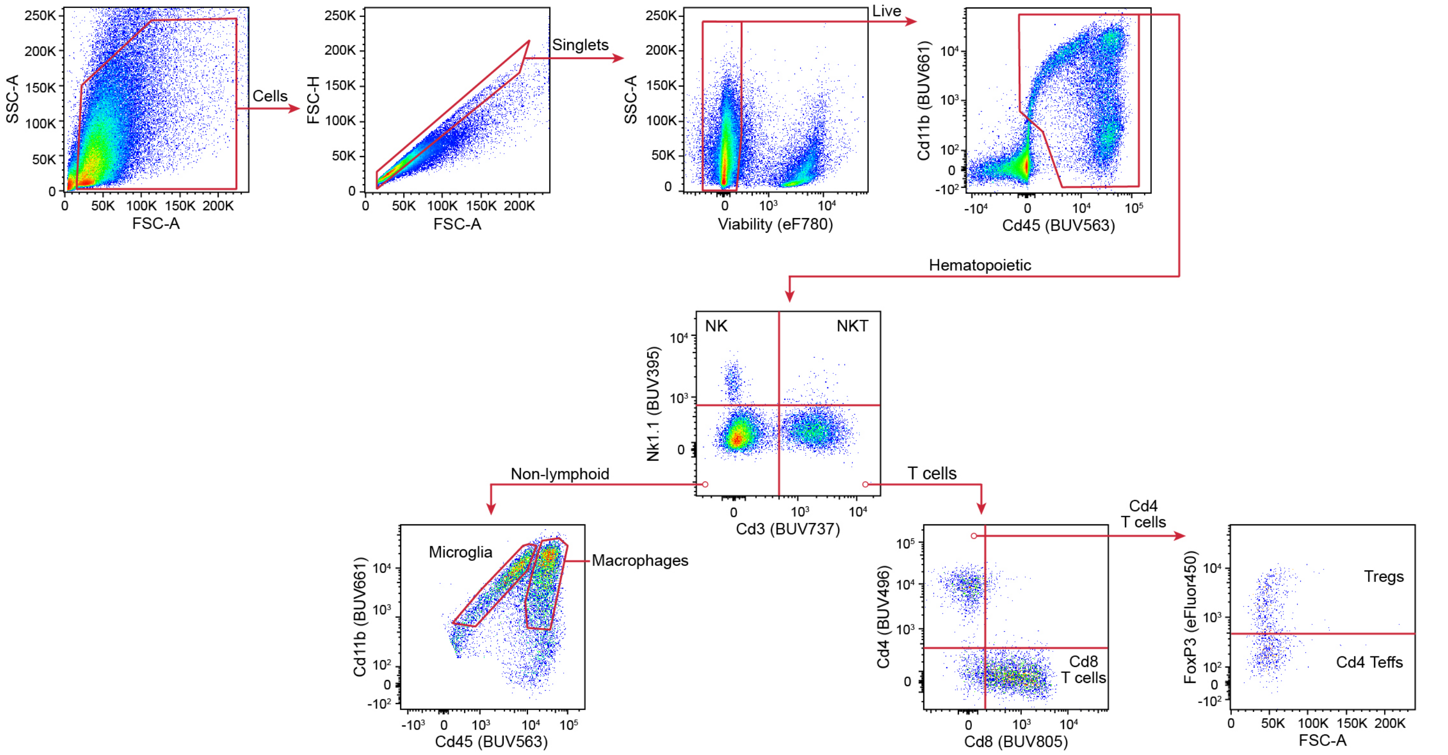

**Supplemental Figure 7. FACS gating strategy**
